# Rfx2 stabilizes Foxj1 binding at chromatin loops to enable multiciliated cell gene expression

**DOI:** 10.1101/085571

**Authors:** Ian K Quigley, Chris Kintner

## Abstract

Cooperative transcription factor binding at cis-regulatory sites in the genome drives robust eukaryotic gene expression, and many such sites must be coordinated to produce coherent transcriptional programs. The transcriptional program leading to motile cilia formation requires members of the DNA-binding forkhead (Fox) and Rfx transcription factor families and these factors co-localize to cilia gene promoters, but it is not clear how many cilia genes are regulated by these two factors, whether these factors act directly or indirectly, or how these factors act with specificity in the context of a 3-dimensional genome. Here, we use genome-wide approaches to show that cilia genes reside at the boundaries of topological domains and that these areas have low enhancer density. We show that the transcription factors Foxj1 and Rfx2 binding occurs in the promoters of more cilia genes than other known cilia transcription factors and that while Rfx2 binds directly to promoters and enhancers equally, Foxj1 prefers direct binding to enhancers and is stabilized at promoters by Rfx2. Finally, we show that Rfx2 and Foxj1 lie at the anchor endpoints of chromatin loops, suggesting that target genes are activated when Foxj1 bound at distal sites is recruited via a loop created by Rfx2 binding at both sites. We speculate that the primary function of Rfx2 is to stabilize distal enhancers with proximal promoters by operating as a scaffolding factor, bringing key regulatory domains bound by Foxj1 into close physical proximity and enabling coordinated cilia gene expression.

**Author Summary:** The multiciliated cell extends hundreds of motile cilia to produce fluid flow in the airways and other organ systems. The formation of this specialized cell type requires the coordinated expression of hundreds of genes in order to produce all the protein parts motile cilia require. While a relatively small number of transcription factors has been identified that promote gene expression during multiciliate cell differentiation, it is not clear how they work together to coordinate the expression of genes required for multiple motile ciliation. Here, we show that two transcription factors known to drive cilia formation, Foxj1 and Rfx2, play complementary roles wherein Foxj1 activates target genes but tends not to bind near them in the genome, whereas Rfx2 can’t activate target genes by itself but instead acts as a scaffold by localizing Foxj1 to the proper targets. These results suggest not only a mechanism by which complex gene expression is coordinated in multiciliated cells, but also how transcriptional programs in general could be modular and deployed across different cellular contexts with the same basic promoter configuration.

## Introduction

Animal gene expression is typically mediated by cell type specific transcription factors, acting through consensus binding sites present in distal enhancers and proximal promoters. Such factors act by opening chromatin,^1,2^ by facilitating the deposition of histone modifications,^3^ by employing local interactions with other factors,^3,4^ or some combination of these. While the details of how discrete sites in the genome accumulate transcriptional machinery are just coming into focus, the mechanism by which these sites bind to and coordinate gene expression at a given promoter is less well known. Recent work hints that this coordination is mediated by 3-dimensional chromosomal architecture.^5–10^ More specifically, genomic regions with high levels of spatial self-interaction, termed topological domains or topologically-associated domains (TADs), have been proposed to encourage promoter-enhancer interactions or transcriptional activation more generally.

Work on discrete transcriptional programs has demonstrated the integration of multiple transcription factors at individual promoter elements.^11^ The production of specialized cilia in animals requires one such transcriptional program: these structures are deployed in a variety of cell types in different anatomical locations but appear to arise in these different settings using a similar transcriptional architecture.^12,13^ For example, the presence of both Fox and Rfx transcription factors at a handful of cilia gene promoters has been associated with robust gene activation in the mammalian airway and also *Drosophila* neurons.^14,15^ However, cilia are composed of hundreds of gene products, and it is not clear how many of them require cooperation between Fox and Rfx factors at promoters or at distal regulatory elements, to what extent other factors are involved, or if changes in 3-dimensional genome structure accompany their specification.

Here we analyze how Foxj1 and Rfx2 cooperate, both at the level of sequence specificity and chromosomal architecture, within *Xenopus* skin progenitors to promote gene expression during MCC differentiation. First, we use extensive RNAseq to capture a robust transcriptome of MCC genes. Next, we use tethered conformation capture, a variant of HiC, to map the organization of topologically-associated domains within epithelial progenitors. We use ChIPseq of the cohesin component Rad21 to confirm these domains and map the location of active regions (H3K4me3 and H3K27ac) and genes expressed during MCC differentiation to them. We further analyze Foxj1 binding using ChIPseq and find that its binding, along with Rfx2, is a good predictor of MCC gene expression as compared to the binding of other major MCC transcription factors such as Myb and E2f4. We further show that Foxj1 stabilization at genomic sites, especially MCC promoters, frequently requires Rfx2. Finally, by examining 3D chromatin interactions, we show that Rfx2, Foxj1, and the promoters of MCC genes lie at the anchor endpoints of DNA loops, and that these interactions are stronger in progenitors converted entirely into multiciliated cells. We propose a model in which dimerization of Rfx proteins recruits or stabilizes distal enhancers to promoters as a scaffolding factor, enabling distantly bound factors such as Foxj1 to promote stable transcription of cilia genes during terminal MCC differentiation.

## Results

### Identification of a discrete transcriptome for MCCs

The MCC is a major cell type in the *Xenopus* larval skin that forms in the early embryo along with mucus-secreting cells and proton-secreting ionocytes. The differentiation of these cell types can be readily analyzed using RNAseq^16^ analysis on skin progenitors isolated away from the embryo. To identify gene expression associated with distinct differentiation programs, we analyzed isolated progenitors where cell fate was altered in predictable way. Blocking Notch in these progenitors (by expressing a DNA binding mutant of suppressor of hairless, dbm^17^, labeled here as “Notch-”) leads to a marked increase in the number of MCCs and ionocytes that arise within an inner layer and move into the outer epithelium (Figure 1B, C). Activating Notch, (by expressing the intracellular domain of Notch, icd^18^, labeled here as “Notch+”) completely suppresses MCC and ionocyte formation. Ectopic expression of Multicilin^19^ in skin progenitors along with active Notch completely rescues MCC differentiation, and moreover, converts most skin progenitors into MCCs, including those in the outer layer normally fated to become mucus-secreting cells. Inhibiting Multicilin activity using a dominant-negative mutation (termed dnMulticilin or dnMcidas^19^) blocks the formation of MCCs, but spares other cell types such as ionocytes. Finally, for an additional comparison, we ectopically expressed Foxi1 in these progenitors, a transcription factor that promotes ionocyte differentiation alone. Epithelial progenitors were isolated from injected embryos at stage 10, and subjected to RNAseq analysis as the various differentiation programs unfolded (Fig. 1C). To maximize the cell-specific signal in these data sets, we compared conditions in which the change in cell number was great: for example, progenitors with activated Notch (using Notch-icd) have virtually no MCCs, whereas progenitors co-injected with activated Notch and Multicilin have large numbers without a confounding ionocyte population. MCC and ionocyte number is also markedly increased by blocking Notch signaling (using dbm), whereas progenitors co-injected with dbm and dnMulticilin have virtually no MCCs but still have increased ionocytes. A complete comparison of the effects of these manipulations is shown in S1 Table. Clustering of genes with the greatest variance (see RNAseq informatics section in Methods) in these extensive RNAseq data sets revealed that the largest cluster of genes by far were those that changed in association with MCC differentiation (Fig. 1F; this group of conditions is detailed in S1 Table and shown in S1 Figure; the cluster of genes themselves is in S2 Table). By comparison, this cluster of gene associated with MCC differentiation dwarfed that associated with ionocyte differentiation (induced with Foxi1 as a control, Fig. 1F, S1A Figure, S3 Table), with Notch signaling (Fig. 1F, S1B Figure, S4 Table), or with embryonic age (Fig. 1F, S1C-D Figure, S5-S6 Tables). These data emphasize the relatively large number of genes coordinately upregulated during the formation of this specialized epithelial cell type.

**Figure 1:**
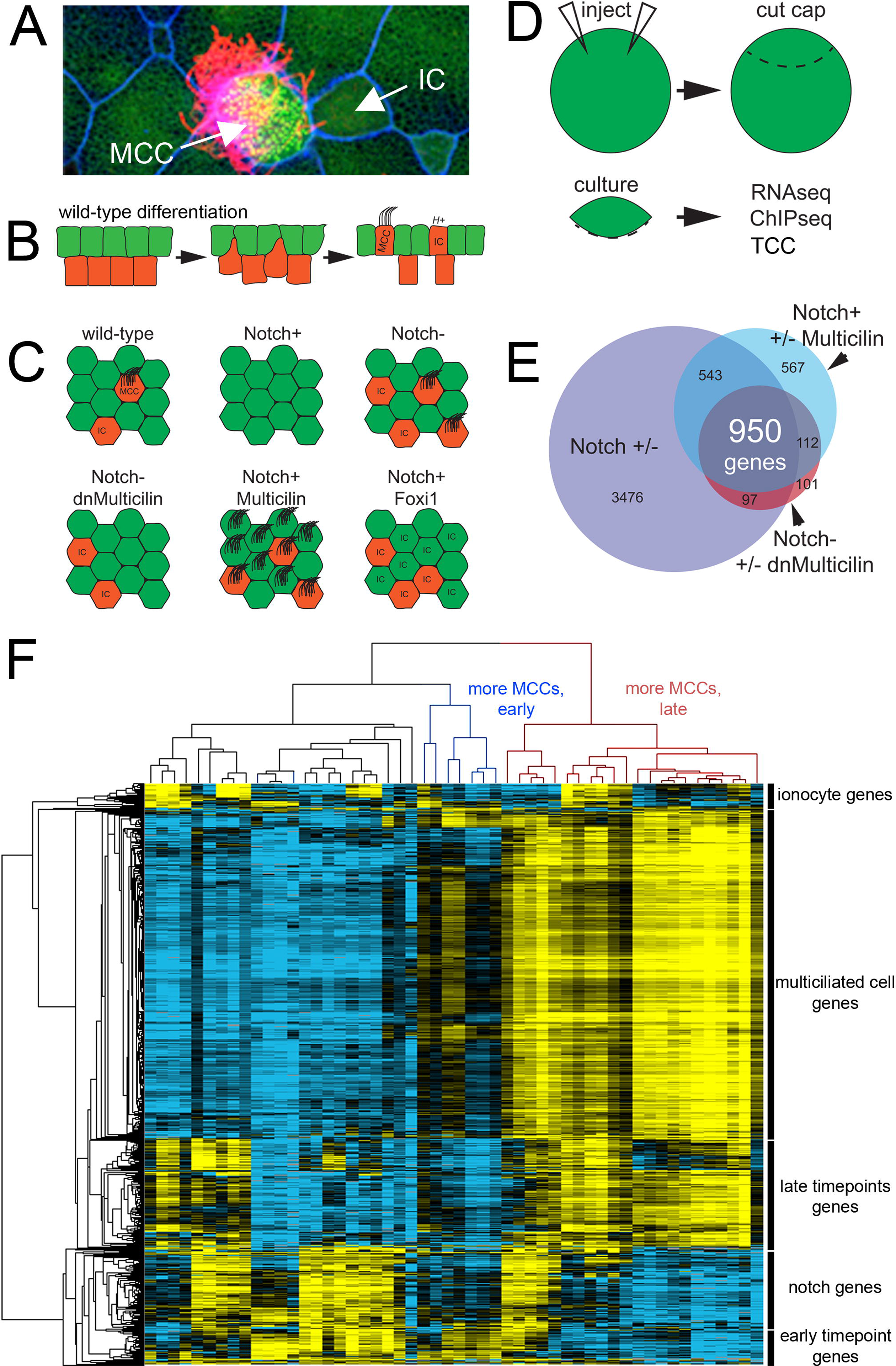
Identification of a MCC transcriptome. (**A**) Confocal image of *X. laevis* skin showing a multiciliated cell (MCC), ionocytes (IC), and outer cells (green cells, unlabeled). **(B)** Differentiation of X. laevis skin. Multipotent progenitors are specified to become MCCs or ionocytes in the inner layer (red cells) by Notch signaling; they then intercalate into the layer of outer cells (green cells). **(C)** Diagrams illustrating how the numbers of MCCs and ICs in the skin change when Notch, Multicilin and/or Foxi1 activity is manipulated. **(D)** Schematic of the general experimental strategy used to analyze *X. laevis* epidermal progenitors (“cap”), after manipulating gene expression using RNA injection. **(E)** Venn diagram using multiple RNAseq experiments to define a core list of genes expressed in MCCs based on an intersectional strategy. **(F)** Heatmap of transcriptional variation across all experimental conditions and timepoints subjected to RNAseq analysis (see Methods for more details). For clarity of display, sample names are omitted here but can be seen in Supplemental figure S1.

We next generated a reliable list of genes upregulated during MCC differentiation based on differential expression (p adjusted for multiple testing < 0.05) between pairs of conditions exhibiting the greatest change in MCC number (Fig. 1B, D: dbm vs. Notch-icd, Notch-icd + Multicilin vs. Notch-icd, and dbm vs dbm + dnMulticilin). A conservative list of 950 genes deriving from the oldest, 9-hour timepoint (813 genes if we collapse the L and S forms based on orthology assignment) that changed in all three comparisons (p<0.05, Fig. 1E) provides a high-confidence collection of the genes upregulated during MCC differentiation, hereafter referred to as “core MCC genes” (S7-9 Tables). To assess this claim, we compared our core MCC gene list to those generated using other approaches, including Foxj1 overexpression in Zebrafish,^20^ FACS sorting of Foxj1^+^ cells from mammalian airway epithelia,^21^ or knockdown of Rfx2 in *Xenopus.*^22^ These gene lists, as well as our own, were plotted using our data based on their level of expression (using normalized counts) versus the fold-change that occurred in our RNAseq analysis as MCC number increased. In each case, the published lists lacked a large fraction of genes on our list (red dots in S2B-D Figure, also see S11-13 Tables for a comparison of overlap between these sets), and the genes unique to our list for each comparison were strongly enriched for cilia GO terms (S3A,C,E Figure). Conversely, the published list included a large number of genes that did not increase with more MCCs in our data, and thus absent from our list (grey dots in S2B-D Figure), but these genes were either not strongly enriched for any GO terms or enriched for more general terms (S3B,D,F Figure e.g., “biological process”). Altogether, these comparisons suggest that the core MCC list more accurately identifies genes upregulated during MCC differentiation, and thus serves as a solid foundation for the subsequent analysis described below.

### 3D chromosomal architecture suggests that MCC genes are enriched at TAD boundaries

We next related the MCC core genes to the 3-dimensional genome, in particular to areas of high self-interaction known as topologically-associated domains (TADs). To obtain 3D chromosomal interactions across the entire genome, we performed tethered conformation capture (TCC)^23^ on epithelial progenitors isolated from *X. laevis* embryos, analyzing both wild-type tissue (unmanipulated progenitors containing a mixture of outer cells, ionocytes, and multiciliated cells), and after inducing MCC differentiation using Multicilin (resulting in progenitors consisting of MCCs). Most of the interactions in our data sets were within chromosomes (92%-87% of totals, respectively, S4 Figure), indicating a low level of interchromosomal interactions (high levels of these are thought to reflect spurious interactions^24^). The active and inactive compartments in the wild-type and Multicilin-injected samples were compared by computing Pearson’s correlation matrices^25^ (Fig. 2A). This analysis failed to detect loci with a negative correlation coefficient and very few with correlation coefficients below 0.5 (268 bins out of a possible 50,030, S14 Table), suggesting that while these two tissues are significantly different in their transcriptional profiles (Fig. 1), there was very little in the way of large-scale changes, such as compartment switching between the two conditions.^7,25^ Genes found in the few regions that were poorly correlated are not implicated in MCC differentiation (S14 Table), while those on our core MCC list, for example, *wdr16*, were highly correlated between the two conditions (Fig. 2A).

**Figure 2:**
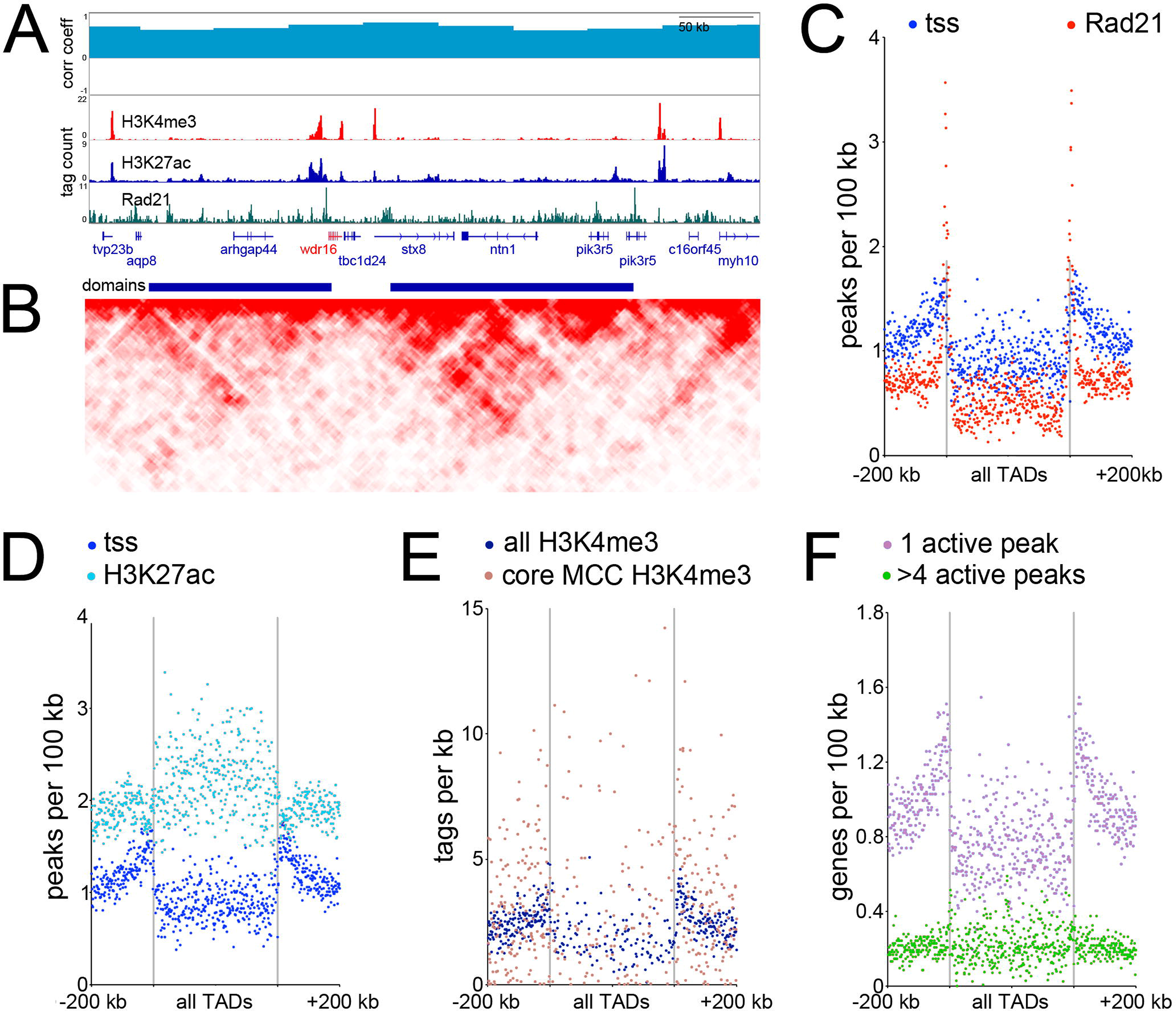
TADs and their genomic features. **(A)** Browser screenshot of the genomic region surrounding *wdr16,* a MCC expressed gene. Top track is correlation coefficient between 3D chromosome conformation data between wild-type ectoderm and ectoderm injected with Multicilin, middle tracks show ChIPseq results as labeled, bottom track is called topological domains. **(B)** Interaction matrix of tethered conformation capture of the same genomic region. High-throughput methods of determining 3-dimensional chromatin structure such as TCC or HiC involve isolating DNA-protein complexes either via dilution (classical HiC), by fixing the proteins to avidin beads (TCC) or *in situ* nuclear fixation (*in situ* HiC), cutting DNA in this folded state with a restriction enzyme at many positions, religating, and sequencing. Restriction sites near loops of DNA will ligate across the loops at some frequency, which can then be used to reconstruct the frequency of contact between two close or distant regions of the genome. Here, regions interacting across the genome more often than a linear model of DNA would predict are shown, with darker red indicating a higher frequency of interactions.^6-9^ **(C-F)** Metagene plots showing the distribution of various features relative to all TADs. All domains are normalized to the same size, the domain region is in the center of each plot, and the two vertical lines denote the domain boundaries. Areas in outer edges of the plot denote flanking genomic regions of some 200 kb. Each quartile of the plot (one quartile upstream, two quartiles inside the domain region itself, and one quartile downstream) is broken into 175 bins, and each dot denotes the measured values of one bin.

At higher resolution, 3D interaction maps reveal TADs, structures thought to facilitate complex transcriptional regulation.^7,9,26,27^ To determine the positions of TADs in the *X. laevis* genome, we pooled the reads from the two TCC experiments and calculated a directionality index,^26^ resulting in 7,249 domains (Fig. 2A,B). As TAD boundaries are reportedly enriched for CTCF and cohesin, we performed ChIPseq on the cohesin component Rad21^26,28^ (Fig. 2A, S5 Table) and found striking enrichment of this protein at our called domain boundaries, providing independent validation that they were called accurately (Fig. 2A,C). We also found Rad21 peaks not associated with TADs (Fig. 2A, ~81% of total peaks); reports by others suggest that both Rad21 and CTCF are frequently outside TADs and are most likely to mark TAD boundaries when cobound.^29^ We annotated the positions of transcriptional start sites (TSS’s) relative to TADs and found enrichment at the boundaries (Fig. 2C), in agreement with previous work on mammalian genomes.^26^ We further performed ChIPseq on two histone modifications: H3K4me3, a mark associated with actively-transcribed promoters (Fig. 2A, S6 Figure), and H3K27ac, a mark associated with active promoters and enhancers, including superenhancers^30^ (Fig. 2A, S7 Figure). The H3K4me3 ChIPseq (Fig. 2A,D, S6 Figure) exhibited broadly similar nucleotide frequencies and peak shapes as other vertebrate promoters^31^ and largely overlapped, within a 2 Kb window, our annotated 5’ ends of *X. laevis* gene models (16,231 out of 24,384 total models; note not all of these genes will be expressed in epithelial progenitors and would thus largely lack H3K4me3 peaks). When these features were overlaid onto the TAD map, the promoter regions, as defined by both annotated TSS’s and H3K4me3 peaks, tended to lie close to the TAD boundaries (Fig. 2C, D), while enhancers were depleted at boundaries and instead enriched in the middle of the domains themselves (Fig. 2D). We localized our core MCC genes onto the TAD map using H3K4me3 binding at their promoters (Fig. 2E) or the positions of the core MCC TSS’s themselves (S8 Figure), finding that, like all genes, core MCC genes were enriched at TAD boundaries, rather than located within domains.

### MCC genes reside in enhancer-sparse regions

The enrichment of MCC genes at TAD boundaries and far from the bulk of enhancers led us to examine how many enhancers were near these genes in greater detail. A single collection of all peaks with active histone marks (H3K4me3 and H3K27ac) was generated and assigned to genes with the closest TSS (S15 Table), based on the assumption that this approach estimates local enhancer density rather than attempting to predict targets of every enhancer. Using this approach, we found an average of 2.84 active peaks closer to a given gene than any other gene across the entire genome. Genes with the highest number of flanking active peaks (> 4) tended to be TFs based on GO term functional analysis (“transcription factor activity”, p value 1.02E-20 for genes with more than 4 flanking active peaks, S15 Table). To confirm this, we checked known TFs (*Xenopus* paralogs of 1,692 human TFs^32^) and found they had 4.04 active flanking peaks on average and were positioned inside TADs, where enhancer density was highest (S9 Figure). Conversely, genes with a single nearby active peak were enriched in GO terms such as “protein transport” (p value=1.7E-6), and “ribosome”, (p value=6.1E-5). Known housekeeping genes (*Xenopus* paralogs of 3,804 human genes^33^) contained 2.40 active flanking peaks on average, and these genes were also enriched at TAD boundaries rather than within them (S9 Figure). Finally, in a general survey, all genes with few enhancers largely resided at the boundaries, while genes with many enhancers were depleted at boundaries but instead located within domains themselves (Fig. 2F). These results are consistent with the idea that housekeeping genes are located at TAD boundaries where enhancer density is low, while developmental regulated genes, such as TFs, are located in areas of high enhancer density within TADs. In this light, it is striking that the MCC-expressed genes averaged few peaks per gene (2.42) and were enriched at TAD boundaries (Fig. 2E), similar to housekeeping genes. Thus, these data indicate that MCC genes, although expressed in a cell type-specific manner, resemble housekeeping genes, raising the question of how these genes become activated in this configuration.

### TF binding motifs associated with MCC promoters

Previous studies have found that proximal regulatory regions of genes expressed in motile ciliated cells are enriched in Rfx binding sites^34,35^ and that in *Xenopus*, Rfx2 binding is found extensively in association with cilia genes.^22^ Consistent with these observations, the binding site for the Rfx proteins was by far, the most enriched motif identified when sequences (+/-100 bases) around the TSS of the core MCC genes (verified by H3K4me3, S6 Figure) were examined for overrepresented motifs using an unbiased approach (Fig. 3A).^3^ We also detected a similar, striking enrichment of Rfx motifs in the promoters of MCC gene orthologs in human, mouse, zebrafish, fruit fly, and sea starlet (Fig. 3B), despite the fact that the core promoter sequences vary substantially across vertebrate and ecdysozoan lineages^36–38^ and cnidarian promoters as well (S10 Figure). By contrast, binding motifs for other factors involved in MCC differentiation, including Forkhead, E2f, and Myb, were also enriched at MCC gene promoters but this enrichment was less impressive, especially at MCC genes in other species (Fig. 3A, S11 Figure).^20,22,39–41^ These data support that idea that Rfx factors directly binding at the promoters of the MCC genes is deeply conserved, but raises the possibility that other factors are recruited to these promoters indirectly.

**Figure 3.**
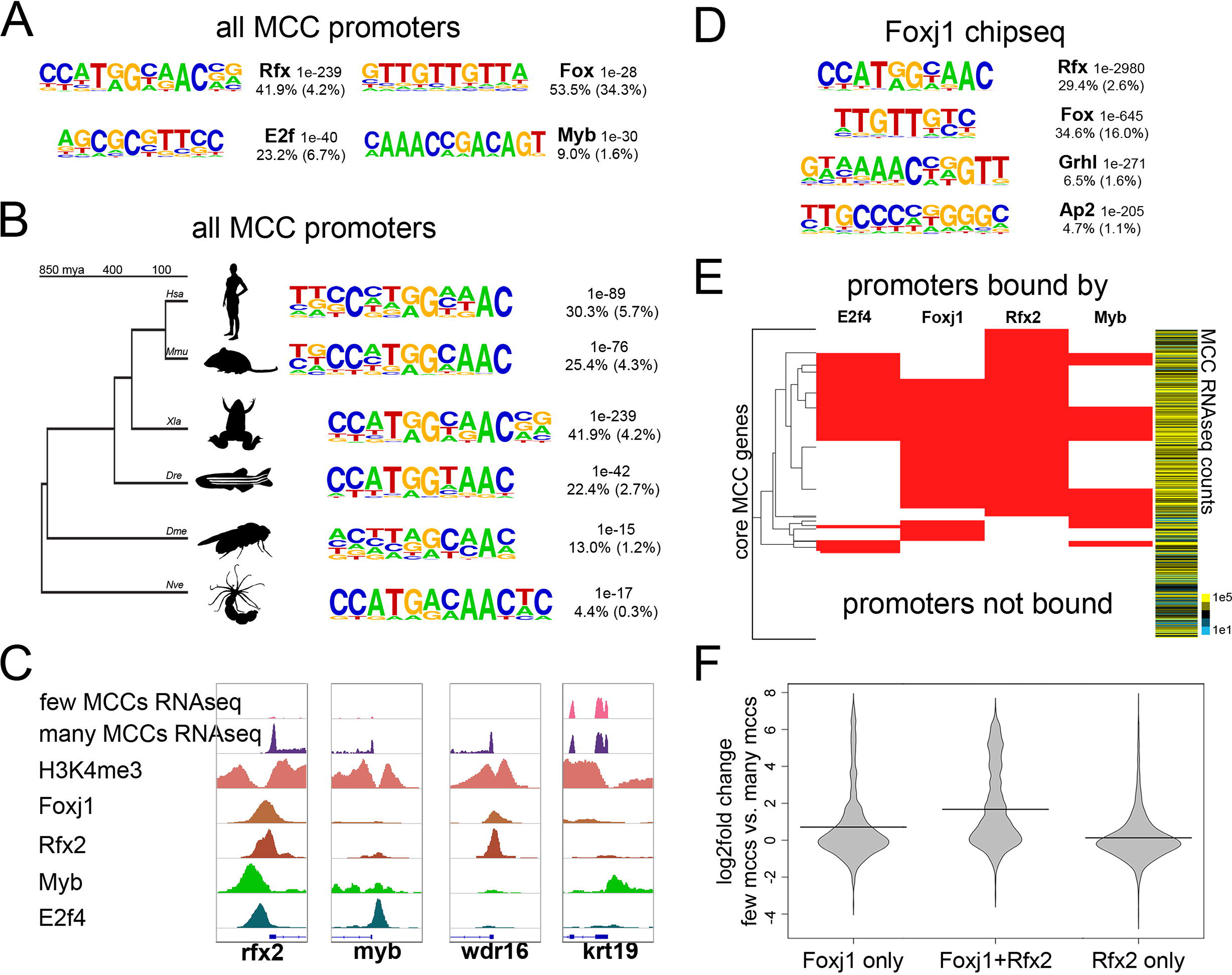
Transcription factor motifs and binding in MCCs. **(A)** Top *de novo* motifs identified in all MCC promoters in *X. laevis* along with the transcription factor family that best matches the motif, a p-value determined by the cumulative hypergeometric distribution, the frequency of the motif in the promoters analyzed, and the background frequency of the motif in all promoters. **(B)** The top *de novo* motif as in (A) that were found in promoters of all MCC paralogs in the indicated species. *Hsa, Homo sapiens; Mmu, Mus musculus; Xla, Xenopus laevis; Dre, Danio rerio; Dme, Drosophila melanogaster; Nve, Nematostella vectensis.* **(C)** Example screenshots around the promoters of three genes that were upregulated during MCC differentiation and one that was not (*krt19*) indicating H3K4me3 or transcription factor binding by ChIPseq. These promoters were bound by various combinations of Foxj1, Rfx2, Myb, and E2f4 as shown. **(D)** Shown are the top *de novo* motifs that are associated with all called Foxj1 ChIPseq peaks. **(E)** Shown are all core 950 MCC promoters and the combinations of E2f4, Foxj1, Rfx2, and Myb bound to each. Heatmap to the right indicates normalized expression counts in manipulations producing many MCCs (epithelial progenitors injected with Notch-icd compared to those injected with Notch-icd and Multicilin). **(F)** Shown is the change in expression between epithelial progenitors with few MCCs (injected with Notch-icd) versus progenitors with many MCCs (injected with Notch-icd and Multicilin) driven by all promoters bound by the indicated factors.

### Transcription factor binding to MCC genes

To gain further insight into the TF binding required for MCC gene expression, we examined the binding targets of TFs recognizing the motifs enriched in MCC promoters: Rfx, Fox, E2f, and Myb. We interrogated recently published *X. laevis* ChiPseq data for Rfx2^22^ and E2f4^40^ and generated ChIPseq data for two additional candidate factors, Myb and Foxj1, using *X. laevis* epithelial progenitors manipulated to increase the number of motile cilia (see Methods, Fig. 3C). We mapped reads from all datasets to the *X. laevis* genome (v9.1),^42^ called peaks against input background, and subjected all peaks to *de novo* motif discovery. One of the top motifs identified in each set corresponded to the motif recognized by the factor immunoprecipitated, suggesting that these family members often bound the appropriate motif directly (Fig. 3D, S12 Figure). However, this analysis also suggests possible co-binding; Foxj1 peaks at MCC genes were highly enriched for Rfx motifs, for example, whereas we saw no such enrichment across all promoters or enhancers (Fig. 3D, S6, S7 Figures).

We exploited the extensive binding data at promoters described above to assess which combination of TFs might account for the upregulation of MCC gene expression (Fig. 3E). We found striking heterogeneity in the number of promoters bound by any one factor; likewise, we found heterogeneity in how often different combinations of factors bound to core MCC promoters. For example, Rfx2 bound a majority of these promoters (579 out of 693 MCC promoters bound by any of these factors), and other factors in this study rarely bound MCC promoters in its absence (114 out of 693, note extensive Rfx2 binding in Fig. 3E). The largest overlap between two factors occurred between Foxj1 and Rfx2 (400), the next between E2f4 and Rfx2 (271), and finally the smallest overlap occurred between Myb and Rfx2 (230). A large fraction of core MCC promoters were not bound by any of these factors (257, or ~27%); however, we note that ~17% of MCC promoters were H3K4me3- (159/950), suggesting that their annotated TSS’s were incorrect.^43^ Manual inspection of a subset of these revealed unannotated 5’ exons, and thus TSS’s, outside of our 2kb TSS window based on Mayball annotations (S13 Figure). These misannotations likely led to an underestimation of TF binding at the promoters of the core MCC gene list.

We asked whether binding of MCC transcription factors at core MCC promoters was associated with increased transcription, and examined fold-changes of target genes between conditions repressing MCCs (Notch-icd) and conditions promoting MCCs (Notch-icd and Multicilin)(Fig. 3F) (We did not examine enhancers with this approach: enhancers are more likely influence transcription of nearby rather than distal genes,^3^ but they often skip over the nearest TSS,^44^ making target promoter prediction difficult). While Rfx2 alone bound many promoters across the genome, few of these genes exhibited increased expression in MCCs; neither did genes whose promoters were bound by E2f4 alone, Myb alone, or either co-bound with Rfx2 (Fig. 3F, S14B, S15B Figures). By contrast, Foxj1 modestly increased expression of genes when bound to promoters alone and strongly increased expression when co-bound to promoters with Rfx2 (Fig. 3F). While known to be involved in MCC differentiation,^41^ Myb did not bind to many MCC promoters (Fig. 3E), did not have a motif in MCC promoters nearly as often as Rfx or Forkhead proteins (Fig. 3A), and was not nearly as associated with MCC gene transcription in promoters it did bind to (S14 Figure), suggesting Myb plays a more minor role in the differentiation of these cells. Other factors may also be involved, such as TP73; however, TP73 is not reported to strongly prefer MCC promoters (~3% of putative MCC genes in mouse),^45^ nor was the TP73 consensus motif strongly enriched in *X. laevis* MCC promoters (Fig. 3A), suggesting it may regulate more potent drivers of MCC genes such as Foxj1 rather than large numbers of MCC genes directly. Taken together, these genome-wide analyses largely support the current model that binding of both Foxj1 and a Rfx factor at a given promoter leads to potent activation of gene expression in MCCs while other combinations or individual factors were much less likely to do so.

### Rfx2 influences Foxj1 binding

The coincidence of Foxj1 and Rfx2 binding at MCC genes was evaluated further by examining this binding in detail across the genome. We first estimated whether co-binding might occur simply by chance by evaluating the level of random overlap within areas of open chromatin (n=1000 iterations), as defined by H3K4me3 and H3K27ac-positive regions. While random overlap of open chromatin predicts co-binding in 2208 peaks (SD 37), actual overlap between Rfx2 and Foxj1 binding occurred significantly more often (3505, p < 0.001, see Methods). We next evaluated where Foxj1 binding occurs in the genome in relation to Rfx2 binding. Foxj1 was rarely bound at promoters where Rfx2 binding was absent (~10%, 1203/11800 peaks, Fig 4A), but Foxj1 was often at promoters where Rfx2 binding also occurred (~42%, or 1487/3505 peaks). To determine if Rfx2/Foxj1 co-binding at promoters happens more often than by chance, we found that a random collection of Foxj1 and Rfx2 peaks (equal to the number of co-binding peaks, half Foxj1, half Rfx2) bound to promoters less often (~36%, or 1274 peaks, SD 27), suggesting that when cobound with Rfx2, Foxj1 was more likely to prefer promoters (p < 0.001, see Methods). Taken together, these data suggest that Foxj1 binding is strongly influenced by the present of Rfx2 binding, particularly at promoters.

**Figure 4:**
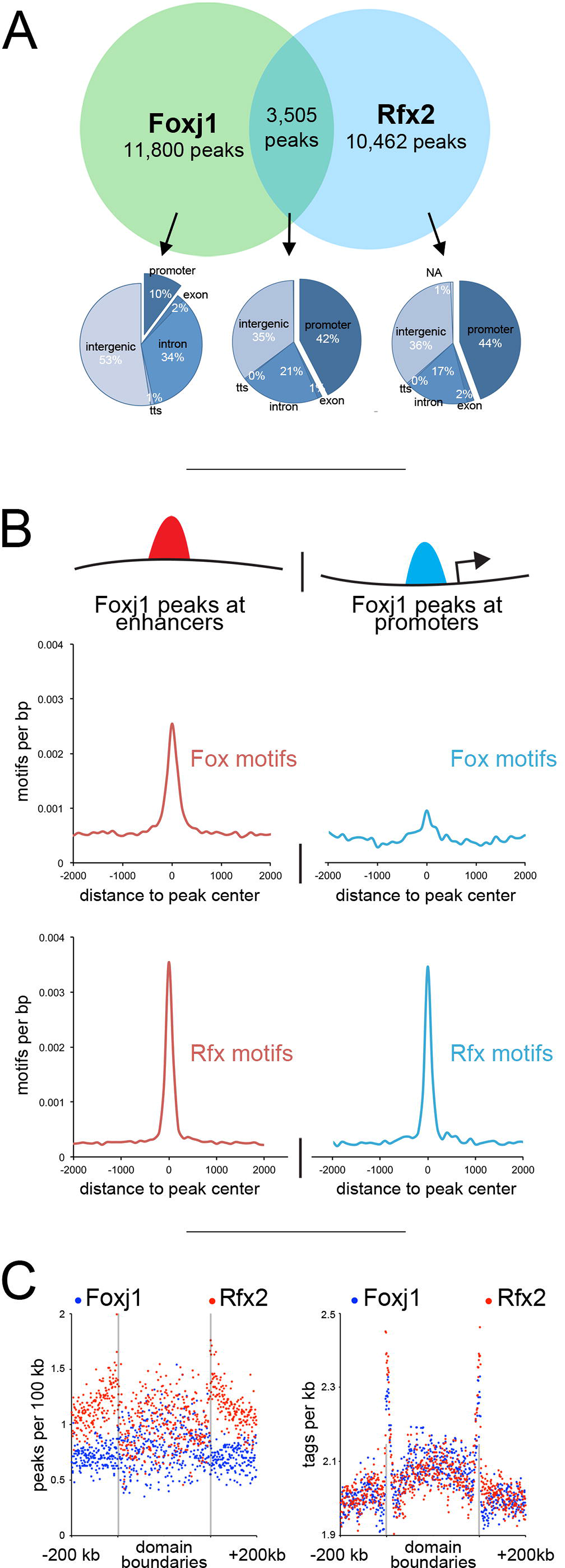
Interactions between Foxj1 and Rfx2. **(A)** Venn diagram at the top represents all peaks (fixed width) bound by Foxj1 and Rfx2 identified with ChIPseq, in terms of their overlap. The pie charts shown below illustrate where Foxj1 and Rfx2 binding occurs in relation to annotated genomic features depending on whether they bind together or alone. **(B)** Foxj1 ChIPseq peaks were subdivided into those located near promoters (< 1 kb to the TSS) or at more distal sites (> 1 kb to the TSS) and then analyzed for the frequency of Rfx or Fox binding motifs. **(C)** Shown is the distribution of Rfx2 and Foxj1 peaks relative to TAD boundaries (left panel), and the distribution of Rfx2 and Foxj1 sequencing tags relative to TAD boundaries (right panel).

We next compared the peaks of Foxj1 and Rfx2 binding to the location of TADs within the *Xenopus* genome. As a factor more likely to be found at distal sites, Foxj1 peaks were enriched inside domains (much like H3K27ac, Fig. 2D), as compared to Rfx2 peaks (Fig. 4C), which were enriched at their boundaries (similar to promoters, Fig. 2C,D). When we examined the numbers of sequencing tags immunoprecipitated from these experiments, however, we saw a striking enrichment for both Rfx2 and Foxj1 at domain boundaries (Fig. 4C), suggesting that while there were fewer Foxj1 peaks at TAD boundaries, those present were exceptionally strong. These results further indicate that Foxj1 binding at MCC genes located at TAD boundaries is associated with the presence of Rfx2 binding.

### Rfx2 stabilizes Foxj1 on DNA

The simplest model to explain co-binding of Foxj1 and Rfx2 at MCC promoters is based on the presence of neighboring direct binding sites with appropriate consensus motifs, as shown in a handful of *Drosophila* cilia genes.^15^ However, when we examined the enrichment of Forkhead and Rfx motifs in the promoters and distal sites bound by Foxj1 (Fig. 4B, also see S16 Figure), we found that Rfx motifs were strongly present to an equal degree in promoters and distal sites, but that Forkhead motifs were depleted in promoters relative to distal sites. Moreover, when we performed *de novo* motif analysis on Foxj1 peaks not co-bound with Rfx2, we again saw robust enrichment of Rfx motifs, suggesting other Rfx proteins may also co-bind with Foxj1 (S17 Figure). Finally, despite Foxj1’s known conserved role in motile cilia formation,^20,35,39,46,47^ Forkhead motifs were not particularly conserved in the promoters of MCC genes across vertebrate lineages (S11 Figure). Thus these findings suggest that the sequences required for the recruitment of Foxj1 to these promoters is dispensable in many cases, even if Foxj1 itself is not.

As there was a strong overlap between Foxj1 and Rfx2 peaks, and few Fox motifs at Foxj1-bound promoters (Fig. 4B), we asked if Rfx2 might stabilize Foxj1 binding at these positions. Moreover, as many Foxj1 peaks were within TADs (Fig. 4C), we speculated that Foxj1 at distal sites might be recruited to promoters, possibly at the TAD boundaries, in chromatin loops and that Rfx2 might facilitate this recruitment. To directly test the possibility that Rfx2 stabilized Foxj1 binding, we knocked down Rfx2 with a well-characterized morpholino^22,48^ and performed ChIPseq on Foxj1 (Fig. 5). Out of 15,305 Foxj1 peaks identified in control tissue, some 6,360 were reduced at least 3-fold (~42%), and 2,084 were reduced 10-fold (~14%) in an Rfx2 knockdown. We saw strong reductions at key promoters, such as the *rfx2* promoter itself (Fig. 5A), while other promoters maintained Foxj1 occupancy (Fig. 5B). We also saw strong reductions in regions with many peaks, such as the superenhancer surrounding *tubb2b* (Fig. 5C); in cases where reduced Foxj1 peaks did not overlap with robust Rfx2 peaks we often saw Rfx motifs (Fig. 5C) and we also found Rfx motifs enriched in Foxj1 peaks not cobound by Rfx2 (S17 Figure), further hinting that other Rfx proteins might be involved. When we looked across all regions bound by Foxj1, we saw a reduction of Foxj1 tags at those positions in the absence of Rfx2 and striking reduction of Foxj1 tags at the promoters of MCC genes (Fig. 5D). By contrast, we saw a slight increase at non-MCC promoters, which may be the result of excess Foxj1 protein binding nonspecifically to regions of open chromatin.^49,50^ While there is a large fraction of Foxj1/Rfx2 cobound sites in promoters that are not upstream of genes in our core MCC list (3025 out of 3505), we found the greatest Foxj1 reduction at core MCC promoters (S18 Figure).

**Figure 5:**
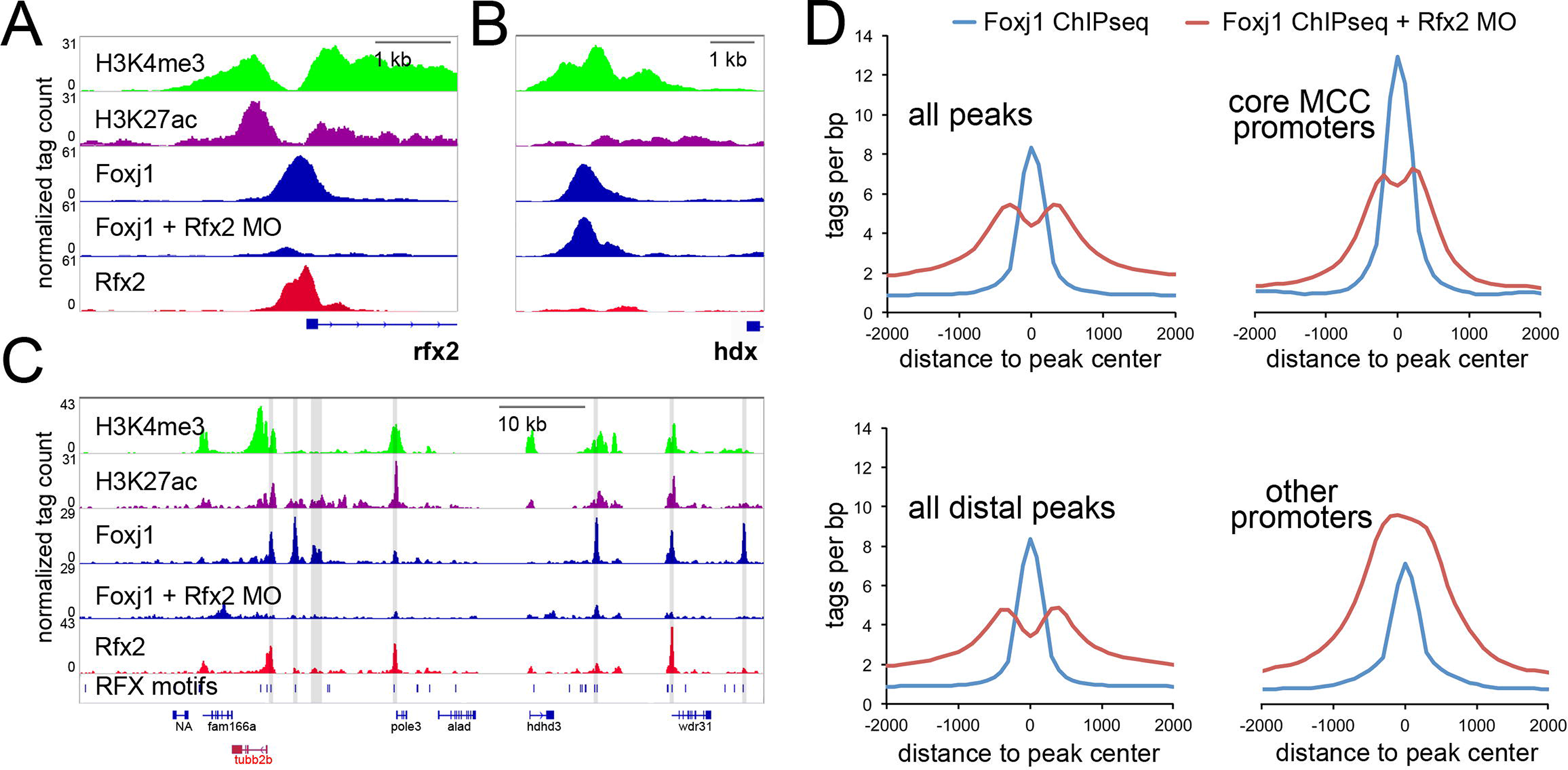
Binding of Foxj1 at promoters is dependent on Rfx2. **(A-B)** Screenshot around the promoter of *X. laevis rfx2* (A) or *hdx* (B) along with the sequence tags obtained in ChIPseq of H3K4me3, H3K27ac, Foxj1, or Rfx2 in wildtype and Foxj1 in Rfx2 morphants (MO). **(C)** Shown is a screenshot of the *X. laevis* genomic region containing the *tubb2b* gene with ChIPseq tracks as in (A). The position of all Rfx motifs is denoted in the bottom track. Foxj1 peaks that are reduced >3-fold in the Rfx2 morphants compared to control are shaded. **(D)** Shown are sequencing tag histograms in peaks as labeled from ChIPseq of Foxj1 in wild-type progenitors or progenitors from Rfx2 morphants.

Rfx2 might stabilize Foxj1 at sites where Foxj1 binds directly to its consensus sequence in DNA, or alternatively, it might stabilize Foxj1 at positions where Foxj1 binds indirectly. To distinguish between these possibilities, we examined changes in Foxj1 binding in peaks occupying positions containing either a strong Forkhead motif but not a Rfx motif (2,646 peaks) or a strong Rfx motif but not a Forkhead motif (2,083 peaks). We found that while tag counts in positions containing a Forkhead motif alone were modestly reduced in the absence of Rfx2, Foxj1 tag counts at positions with an Rfx motif alone were drastically reduced (S18 Figure), suggesting that Rfx2 might have a minor role in stabilizing Foxj1 directly but a much larger role in stabilizing it indirectly.

### Rfx2 stabilizes Foxj1 inside chromatin loops

The increased likelihood of transcription of promoters bound by multiple factors is consistent with general findings by others^3,11^ but does not explain the distribution of Foxj1, Rfx2, and their respective motifs relative to transcriptional start sites. Rather, the results above suggest a model where, in some cases, Rfx proteins bind at both promoters and distal enhancers, and, through dimerization,^51,52^ maneuver the two together to effect transcription. We further speculated that this looping enables Foxj1, bound to enhancers, to localize to promoters, where it was then crosslinked and immunoprecipitated in our assays. This proposed looping would create a protein-DNA complex wherein Foxj1 bound enhancer DNA through a motif but was held nearby its target promoter DNA via Rfx2 proteins: ChIP of a single Foxj1 protein would then pull down both its directly-bound enhancer and its indirectly-bound promoter. To evaluate this model, we first determined all significant interactions in our 3D chromosomal conformation data from wild-type epithelial progenitors using an approach published previously (see Methods, “*TCC significant interactions and TADs*”).^25^ We then interrogated the anchor points of these interactions for co-binding of Rad21, surmising that the anchor points of significant chromosomal interactions would be enriched for known looping machinery.^28,53^ To test the probability of overlap between positions bound by Rad21 and the anchor endpoints of significant interactions, we performed a hypergeometric distribution test, calculating the amount of overlap by chance alone, the amount of enrichment over this amount measured in our data (if any), and the probability of encountering this enrichment.^3,54^ Pairs of Rad21 peaks were found strongly enriched at the endpoints of loop anchors (Rad21 self-interacting loops in Fig. 6A), in agreement with^53^. We then applied this approach to other peaksets, finding strong enrichment for pairs of Rfx2 peaks at the endpoints of looping interactions, consistent with our model that Rfx2 could mediate such interactions. We also saw pairs of Foxj1 enriched at the ends of loops, as well as Rfx2-dependent Foxj1 peak pairs (labeled as “F3”, for peaks reduced by 3-fold or more in the absence of Rfx2). There was also a strong enrichment for pairs of MCC promoters at the endpoints of loops. Finally, we asked if the peaksets enriched at anchor endpoints changed during MCC differentiation, by comparing wild-type interaction data with data obtained with progenitors expressing Multicilin (Fig. 6B). We saw an increase in enrichment of both Foxj1 peaks and Rfx2-dependent Foxj1 peaks at anchor endpoints, and an even stronger enrichment of endpoints with Rfx2-dependent Foxj1 peaks at one end and MCC promoters at the other end (connection between “F3” and “MCC”). Taken together, these data suggest a model wherein Foxj1 peaks bound to distal enhancers are recruited to MCC promoters via Rfx2 dimerization, a process that coordinates or stabilizes chromatin loops (Fig. 6C-D).

**Figure 6:**
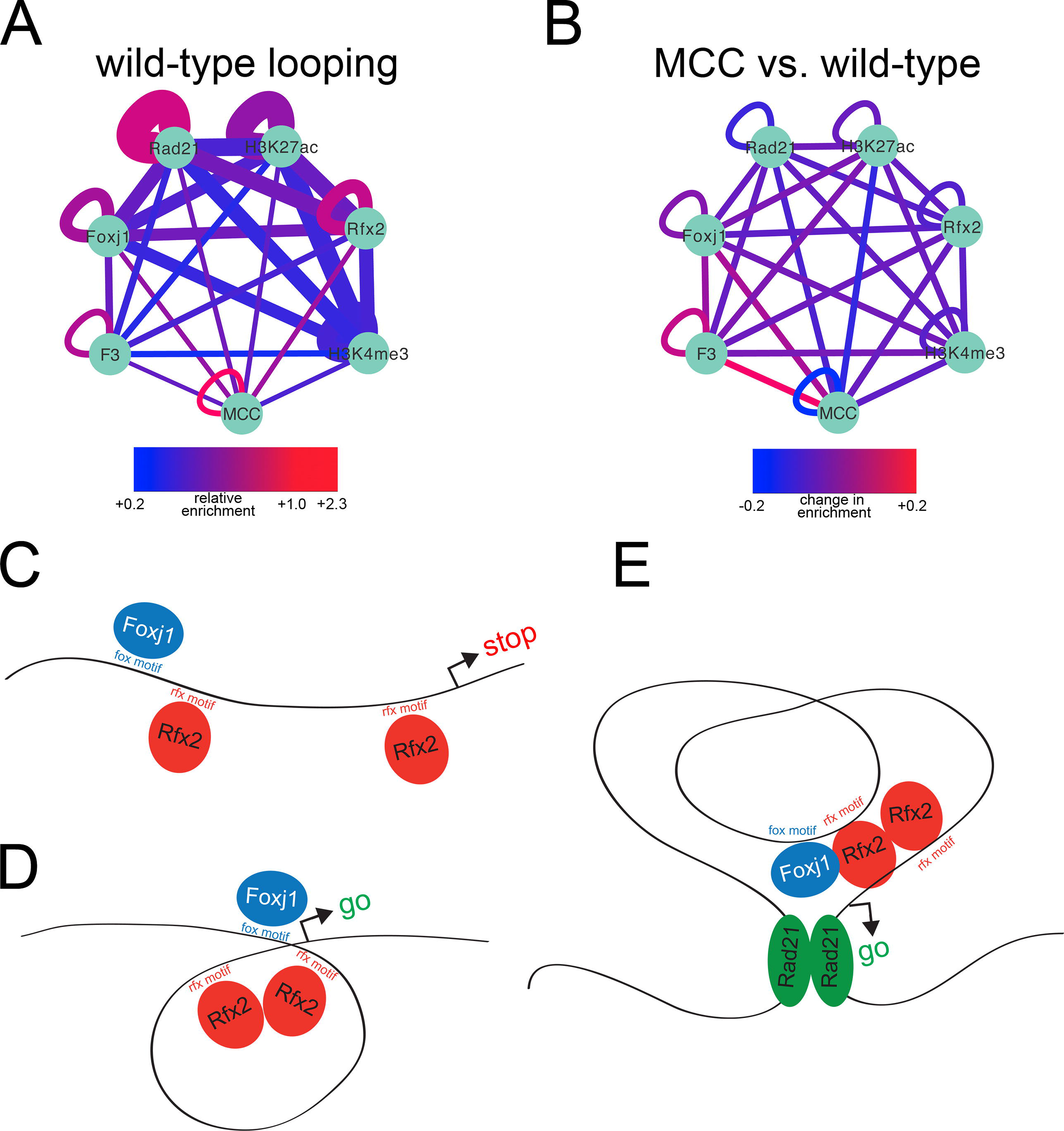
Chromatin loops connect MCC regulatory elements. **(A)** The relative enrichment over expected of histone modifications or transcription factor binding sites at loop anchor points was calculated and visualized in Cytoscape. "Wild-type" tissue is unmanipulated progenitors containing a mixture of outer cells, ionocytes, and multiciliated cells. Expected overlap was determined by hypergeometric distribution; 3D interactions were obtained from wild-type progenitors, and line thickness is inversely proportional to p value (range: 1e-37 to 1e-611, thicker line is lower p value). Nodes are as labeled; "F3" represents the subset of Foxj1 ChIPseq peaks that are reduced 3-fold or greater in Rfx2 knockdowns and "MCC" represents MCC TSS’s. **(B)** 3D interactions were obtained for wild-type progenitors using progenitors injected with Multicilin to increase numbers of MCCs as background (to determine interactions stronger in wildtype tissue) and 3D interactions were also obtained using the reverse (to determine interactions stronger in multiciliated cells). Relative enrichments of histone modifications or transcription factor binding sites were determined for each as in (A) and then compared to one another. Thus, values here depicted by color represent changes in enrichment between the two conditions. **(C,D)** Model of recruitment of Foxj1-bound enhancers to MCC promoters via Rfx2- dimerization. **(E)** Model of how Rfx2-mediated enhancer recruitment operates in the context of TAD boundaries.

## Discussion

TFs required in epithelial progenitors to drive gene expression during MCC differentiation have been identified, but how these factors cooperate at both individual positions in the genome and also within larger chromosomal topologies are largely unknown. Here, using a battery of genomic approaches, many for the first time in *X. laevis,* we show that genes involved in MCC differentiation typically reside at TAD boundaries and are activated by a combination of Foxj1 and Rfx2. Our data suggest a model where Foxj1, often bound directly to flanking enhancers, is recruited to the promoters of MCC genes via Rfx2, which acts as a scaffolding factor (Fig. 6C-E). We speculate that this arrangement facilitates terminal differentiation by maintaining stable activation of gene expression.

We show, using an unbiased approach,^3^ that the MCC core promoters in *Xenopus* and other metazoans are highly enriched for motifs recognized by the Rfx factors, also known as the X-box, in line with a similar analysis of genes differentially expressed in lung tissue from patients with primary ciliary dyskinesia.^55^ While the importance of Rfx binding sites is well established for cilia genes in flies and worms,^12^ their exact role in promoting cilia gene expression in vertebrates is less clear, mainly because the family of Rfx factors has expanded significantly, members of this family appear to act redundantly, and the phenotype associated with a loss of any one family member may differ depending on the species (e.g. Rfx2 in mouse and *Xenopus*).^22,48,56^ In addition, the expanded Rfx family in vertebrates clearly has major roles in enabling gene expression that serves non-cilia functions,^57^ based on single family member mutants (S2 Figure) as well as compound mutants.^34^ The Rfx family also appears to have broad functions based on the high occurrence of Rfx binding motifs within non-coding regions that are conserved within mammalian genomes.^58^ Thus, the analysis of MCC gene promoters suggests that Rfx factors are major players in MCC gene expression but other observations suggest they act by facilitating the action of other TFs. If so, what factors are involved and by what mechanism?

To address this question, we generated ChIPseq data for Myb and Foxj1, two other regulators required for MCC differentiation whose binding preference in epithelial progenitors was unknown. These data, combined with the previously published ChIPseq data for E2f4 and Rfx2,^22,40^ indicate that among the various combinations, the co-binding of both Foxj1 and Rfx2 at a promoter is the best predictor of activated expression during MCC differentiation. By examining the nature of transcription factor co-binding in our data, we uncovered several important features of this synergistic interaction at MCC promoters. First, ChIPseq data indicate that Foxj1 alone preferentially binds to distal sites, but shows a much higher preference for promoters when co-bound with Rfx2. Second, Foxj1 binding at both locations are highly enriched in Rfx motifs, but Foxj1 sites at promoters rarely have strong Forkhead motifs. This may be explained by fixation: we suspect that chromatin immunoprecipitation of individual proteins yields genomic fragments both directly and indirectly bound to that protein due to crosslinking; if so, ChIP of a single Foxj1 molecule might produce fragments from both a distal enhancer and a promoter, and we predict direct binding at fragments with motifs and indirect association at fragments without. Third, Foxj1 binding at MCC promoters and at sites lacking Forkhead motifs is severely disrupted when Rfx2 function is impaired. Taken together, these data indicate that Foxj1 transcriptional activity, especially at MCC promoters and sites with poor motifs, is directed by, and dependent on, Rfx2. These data lead us to propose that Rfx2, perhaps with other Rfx proteins, performs a scaffolding role wherein distal Foxj1-bound enhancers are maneuvered to promoters by Rfx. This scaffolding function may cause looping directly since the Rfx proteins can form homo- and heterodimers with one another,^51,52^ but one can envision more complicated scenarios where Rfx functions within a larger protein complex required for looping and Foxj1 recruitment or Foxj1 binding DNA indirectly through Rfx proteins and dispensing with Fox motifs entirely. While further work will be required to determine if and how the 3D genome is perturbed when Rfx function is lost and how different Rfx proteins might cooperate to affect its conformation, this scaffolding model is consistent with extensive functional data indicating that Foxj1 is the critical limiting factor required for MCC gene expression,^39,59^ explains why Rfx motifs and binding are so frequently encountered at MCC promoters, and is largely consistent with other examples where Rfx and Forkhead factors have been shown to interact to promote gene expression, not only in cells with motile cilia^14,15^ but also in other contexts.^60^ Also consistent with a scaffolding, and not a transcriptional, role, recent work also suggests that Rfx proteins are unable to rescue ciliogenesis in the absence of Foxj1.^45^ Moreover, the scaffolding function of the Rfx proteins proposed here may be more general, enabling enhancer and promoter interactions not only during MCC cell differentiation, but in other contexts as well where this family of proteins is known to act.

Chromatin looping is thought to mediate promoter-enhancer interactions, bringing two or more elements close to one another in topological space.^5,61,62^ Here, TCC data support the presence of such loops maneuvering Foxj1 to MCC genes in that there is enrichment of Foxj1 binding sites of MCC promoters at opposite anchor endpoints of chromatin loops in MCC cells. This finding along, with the observation that MCC genes typically reside at TAD boundaries, may be significant in the context of terminal differentiation. The boundaries that define TADS have relatively stable anchor points, contain a low density of local enhancers, and are highly enriched in transcriptional machinery.^26,53,63,64^ These properties may explain why TAD boundaries also contain a high density of housekeeping genes that are constitutively expressed across most cell types.^64^ Thus, we speculate that positioning of certain MCC genes at TAD boundaries may facilitate stable gene expression associated with terminal differentiation, specifically that associated with the maintenance of the multiple motile cilia in the long-lived MCC.

This arrangement may also provide a modular framework for transcriptional control. While core promoter machinery may impose sequence constraints on GC-rich promoters that are incompatible with AT-rich motifs such as those favored by Foxj1, distal enhancer sequences may be more relaxed.^65^ Thus, distal enhancers may be free to contain a wider range of motifs, and, as long as they also contain flanking Rfx motifs, such elements can be recruited to promoters via Rfx dimerization. As the various Rfx family members may have slightly different motif preferences, one can also speculate that such an arrangement provides for dynamic transcriptional control across tissues and conditions depending on which Rfx proteins are expressed: a distal enhancer in one cell type may get recruited to its target promoter, whereas a different enhancer might get recruited to the same promoter in another cell type.

## Methods

### Embryos, injection, and isolation of epithelial progenitors

*X. laevis* embryos were prepared by in vitro fertilization using standard protocols.^66^ Synthetic, capped mRNA was generated in vitro using DNA templates described below and injected into embryos at the two-cell or four-cell stage (typically 0.1–5.0 ng of RNA per embryo), targeting the four animal quadrants. Pluripotent epidermal progenitors (animal caps) were isolated at stage 9-10 and cultured in 0.5X MMR and harvested at 3, 6, and 9 hours at 22 deg after mid-stage 11. Progenitors injected with RNA encoding HGR-inducible constructs (Multicilin-HGR and Foxi1-HGR) were induced with 1µM dexamethasone at mid-stage 11.

### DNA constructs and knockdowns

DNA templates for synthesis of mRNAs have been described previously for Notch-icd, dbm, Foxi1-HGR, Multicilin-HGR, and dominant negative Multicilin.^18,19,67^ The DNA template for generating an RNA encoding a tagged form of Foxj1 was obtained by subcloning a cDNA encoding Foxj1^39^ into a CS2 vector with a FLAG tag, while that for a tagged form of Myb was generated by PCR amplification of a *myb* cDNA from a stage 17 cDNA library and subcloning into a CS2 vector with a GFP tag. The Rfx2 morpholino^22,48^ was a kind gift of Meii Chung and John Wallingford.

### RNAseq libraries

RNAs were isolated by the proteinase K method followed by phenol-chloroform extractions, lithium precipitation, and treatment with RNase-free DNase and a second series of phenol-chloroform extractions and ethanol precipitation. RNAseq libraries were then constructed (Illumina TruSeq v2) and sequenced on an Illumina platform. Details on specific experiments are in S17 Table, and RNAseq reads are deposited at NCBI (<u>GSE76342</u>).

### RNAseq informatics

Sequenced reads from this study or obtained previously^22,40^ were aligned to the *X. laevis* transcriptome, MayBall version^22^ with RNA-STAR^68^ and then counted with eXpress.^69^ Effective counts from eXpress were then clustered (Cluster 3.0^70^, log transformed, maxval – minval = 6) and visualized with Java Treeview (v1.1.6r, ^71^) to produce heatmaps. DESeq was used to estimate dispersion and test differential expression using rounded effective counts from eXpress.^72^ Changes in expression were visualized in R with beanplot (https://cran.r-project.org/web/packages/beanplot/beanplot.pdf), and to visualize RNAseq reads in a genomic context they were mapped to genome version 9.1 with bwa mem^73^ and loaded as bigWig tracks into the Integrative Genomics Viewer browser.^74^

### Multispecies comparisons

Orthologs/paralogs of MCC genes were obtained from gene assignments in Homologene or by using official gene symbols in *N. vectensis* gene models.^75^ Promoter collections for *H. sapiens, M. musculus, D. rerio,* and *D. melanogaster* were obtained from UCSC. To determine *N. vectensis* promoters, 5’ ESTs were mapped to JGI genome build v1.0 with BLAT.^76^ H3K4me3 ChIPseq reads from *N. vectensis*^77^ were then mapped to v1.0, peaks called, and then overlapped with TSS’s as determined by 5’ ESTs to verify promoters, similar to our approach in confirming *X. laevis promoters.* 5’ ESTs were then aligned to *N. vectensis* gene models (v1.0) to connect transcriptional start sites to gene identity. Motif enrichment on all promoters and MCC ortholog promoters was done with HOMER.^3^ Species divergence times are from Timetree.^78^

### ChIPseq libraries

Because ChlP-grade antibodies are generally not available that recognize *Xenopus* proteins, we tagged Myb and Foxj1 with GFP and FLAG, respectively, and injected mRNAs encoding these constructs into embryos. Myb has functions in stem cell maintenance independent of a role in MCCs,^79^ so to enrich for motile cilia targets, we performed ChIPseq on Myb in progenitors injected with Multicilin-HGR induced at mid-stage 11 (our previous work with E2f4 ChIPseq^40^ also included Multicilin injections, allowing for direct comparison of MCC-enriched TF targets). Moreover, overexpression of Foxj1 promotes ectopic cilia, so samples injected with that construct were also enriched for motile cilia targets by virtue of expressing a tagged Foxj1 construct.

Samples were prepared for ChIP using described methods^40^ with the following modifications: About 250 animal caps for TFs or 100 caps for histone modifications were fixed for 30 min in 1% formaldehyde, and chromatin was sheared on a BioRuptor (30 min; 30 sec on and 2 min off at highest power setting). Tagged proteins with associated chromatin were immunoprecipitated with antibodies directed against GFP (Invitrogen catalog no. A11122, lot no. 1296649) or FLAG (Sigma, cat #F1804). Native proteins were immunoprecipitated with antibodies directed against H3K4me3 (Active Motif, cat #39159; lot #01609004), H3K27ac (Abcam, cat #ab4729, lot #GR71158-2), or rad21 (Abcam, cat #ab992; commercially-available CTCF antibodies target regions not conserved in the *X. laevis* protein). DNA fragments were then polished (New England Biolabs, end repair module), adenylated (New England Biolabs, Klenow fragment 3′–5′ exo- and da-tailing buffer), ligated to standard Illumina indexed adapters (TruSeq version 2), PCR-amplified (New England Biolabs, Phusion or Q5, 16 cycles), and sequenced on an Illumina platform. Details on specific experiments are in S17 Table, and ChIPseq reads are deposited at NCBI (GSE78176).

### ChIPseq informatics

ChIP-seq reads from this study or published previously^22,40^ were mapped to *X. laevis* genome v9.1 with bwa mem,^73^ peaks called with HOMER^3^ using input as background. Peak positions were annotated relative to known exons (Mayball gene models^22^), with promoters defined as being +/- 1Kb around the TSS. Peak sequences were interrogated for *de novo* motif enrichment with HOMER and MCC promoters were clustered (based on if they were bound/not bound) with Cluster 3.0 and visualized with Java Treeview (v1.1.6r). Tags or motifs were counted at peak positions with HOMER and plotted with Excel or R. To determine enrichment statistics, we used the hypergeometric distribution to evaluate the likelihood of peak overlap.^3^ Additionally, as regions of open chromatin occupy a fraction of the entire genome, we also assessed overlap between pairs of factors by determining the distribution of randomly picked open chromatin sites (equal to the number of sites in the first and second TFs in the comparison; open chromatin was estimated by using all H3K4me3 and H3K27ac sites^80^) and repeated this approach 1000 times to get a distribution of overlap between two sets of random regions of open chromatin. We then compared this distribution to the overlap observed in pairs of measured TFs, again restricted to open chromatin (as in Figure 4 and S14, S15 Figures). To determine if subsets of overlapped peaks preferred promoters or other features more than by chance, we employed a similar approach: instead of selecting overlapped peaks, we sampled from all peaks, half from each set, and determined how many peaks form this null set bound to promoters. We repeated this operation 1000 times to obtain the distribution of promoter preference from these random combinations. Finally, when determining differential binding and generating bigwig files for visualization, each experiment was normalized to 10M reads in HOMER-style tag directories to account for differences in sequencing depth.

### Tethered conformation capture

TCC was performed as described^81^ with the following modifications: 100 animal caps were harvested at the 9 hour timepoint and fixed for 30 minutes in 1% formaldehyde. Tissue was thawed, rinsed in PBS, and lysed directly in 250 µl ice-cold wash buffer with SDS added (50 mM Tris.HCl pH=8.0, 50 mM NaCl, 1 mM EDTA, 0.56% SDS) rather than using lysis buffer. A complete protocol is available on request. Details on specific experiments are in S17 Table, and TCC reads are deposited at NCBI (GSE76363).

### TCC informatics

TCC reads were cleared of PCR duplicates, trimmed down to exclude MboI restriction sites (GATC) and mapped to the *X. laevis* genome v9.1 with bwa mem. Topological domain peakfiles, interaction matrices, and metagene plots were generated with HOMER (additional details below). Matrices were visualized with Java TreeView.

### TCC background model

All TADs and significant interactions were called against a background of expected interactions. This background model was constructed using the approach described in ^25^. Briefly, the frequency of proximity-ligated fragments that are connected across a certain distance on a linear chromosome is expected to increase as that distance shrinks. To generate the model, the genome is broken into bins with a sliding-window approach to boost signal, and sequence depth in those regions is normalized to account for mapping errors and biases in restriction enzyme accessibility. Once depth is normalized in all regions, the expected variation in interaction between regions is calculated as a function of distance.

### TCC significant interactions and TADs

As the background model estimates how many interactions between regions are expected given a specific linear distance and sequencing depth, we looked at the number of interactions we actually observed between regions in the TCC data. To determine interactions with some measure of statistical confidence, we used the cumulative binomial distribution, where the number of trials was the total number of reads mapping within regions, success was the expected interaction frequency, and observed successes came directly from read pairs connecting regions from the TCC data itself. We then used the hypergeometric distribution to ask if the anchor points of these interactions overlapped with ChIPseq peaks or other genomic features more than would be expected by chance (Figure 6A) and visualized these values with Cytoscape.^82^ We also determined interactions that were stronger in wild-type or Multicilin-injected epithelial progenitors: after determining significant interactions for one of these conditions, we used TCC reads from the other to score their strength (Figure 6B). TADs were determined by calculating a directionality index with HOMER; we observed similar numbers of TADs with varying resolutions (10-25 kb). All methods in this section were adopted from ^25^.

## Acknowledgements

The authors wish to thank Drs. Chris Benner, Shawn Driscoll, Sven Heinz, and Ronan O’Malley for helpful discussions and technical advice, and Drs. Jesse Dixon and Rhiannon Macrae for comments on the manuscript. This paper is dedicated to the memory of Chung-Ting Chen, a passionate and insightful colleague with a taste for barbecue.

---

**Supplemental figure S1:**
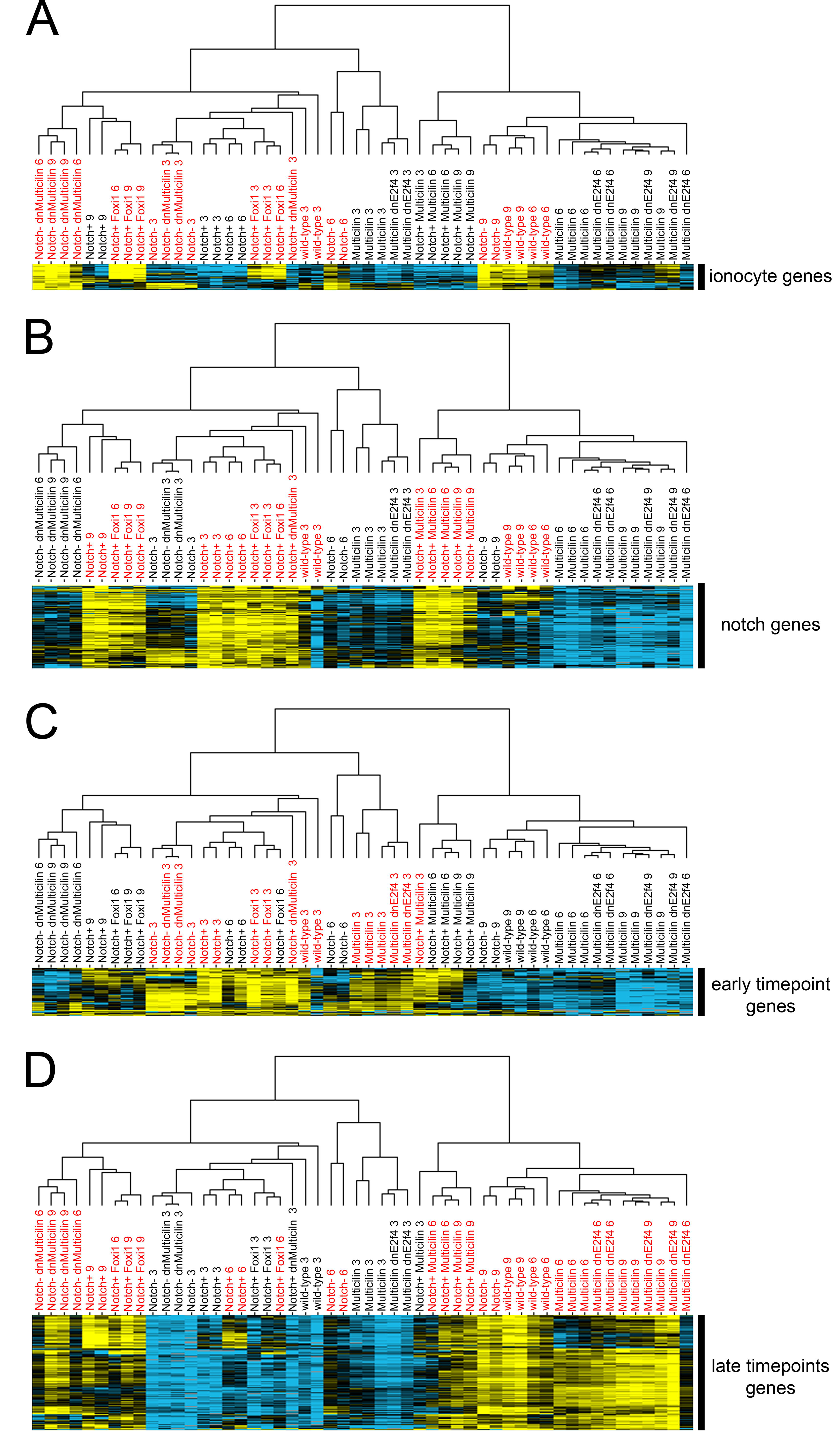
Heatmap of genes associated with the differentiation of ectodermal progenitors. RNAseq analysis was carried out on epithelial progenitors that were isolated from embryos injected with RNA to increase (Notch-icd, labeled as Notch+ (Coffman, Harris, and Kintner 1990; Jennifer L Stubbs et al. 2006)) or decrease (DNA-binding mutant of Suppressor of Hairless or DBM, labeled as Notch- (Deblandre et al. 1999; Jennifer L Stubbs et al. 2006)) Notch signaling, to increase (Multicilin, (J L Stubbs et al. 2012)) or decrease (dominant-negative Multicilin or dnMulticilin, (J L Stubbs et al. 2012) Multicilin activity, to increase (Foxi1, (Quigley, Stubbs, and Kintner 2011)) Foxi1 activity, or to inhibit E2f4 (E2f4ΔCT, (Ma et al. 2014) activity. Samples were collected at 3, 6 or 9 hour timepoints, corresponding to the equivalent of stages 13, 16, and 18, respectively, and hierarchical clustering performed on both samples and expression. **(A-D)** Shown are groups of genes clustered by similarity of expression. Each group shows increased expression corresponding to subsets of experimental treatment. In each case, these treatments are in red (e.g., in (A), Notch-, Notch- Foxi1+, Notch- dnMulticilin, and wildtype treatments all have more ionocytes than Notch+ or Multicilin treatments). **(A)** Genes clustered by similarity of expression found to be increased in treatments promoting ionocytes (also see Table S2). **(B)** Genes clustered by similarity of expression found to be increased in treatments with more Notch signaling (also see Table S3). **(C)** Genes clustered by similarity of expression found to be increased in treatments harvested at the earliest timepoint (3 hours after mid-stage 11, or roughly stage 13; also see Table S4). **(D)** Genes clustered by similarity of expression found to be increased in treatments harvested at the later timepoints ((6 and 9 hours after mid-stage 11, or roughly stages 16 and 18; also see Table S5). This group contains genes associated with the differentiation of goblet and small secretory cells.

**Supplemental figure S2:**
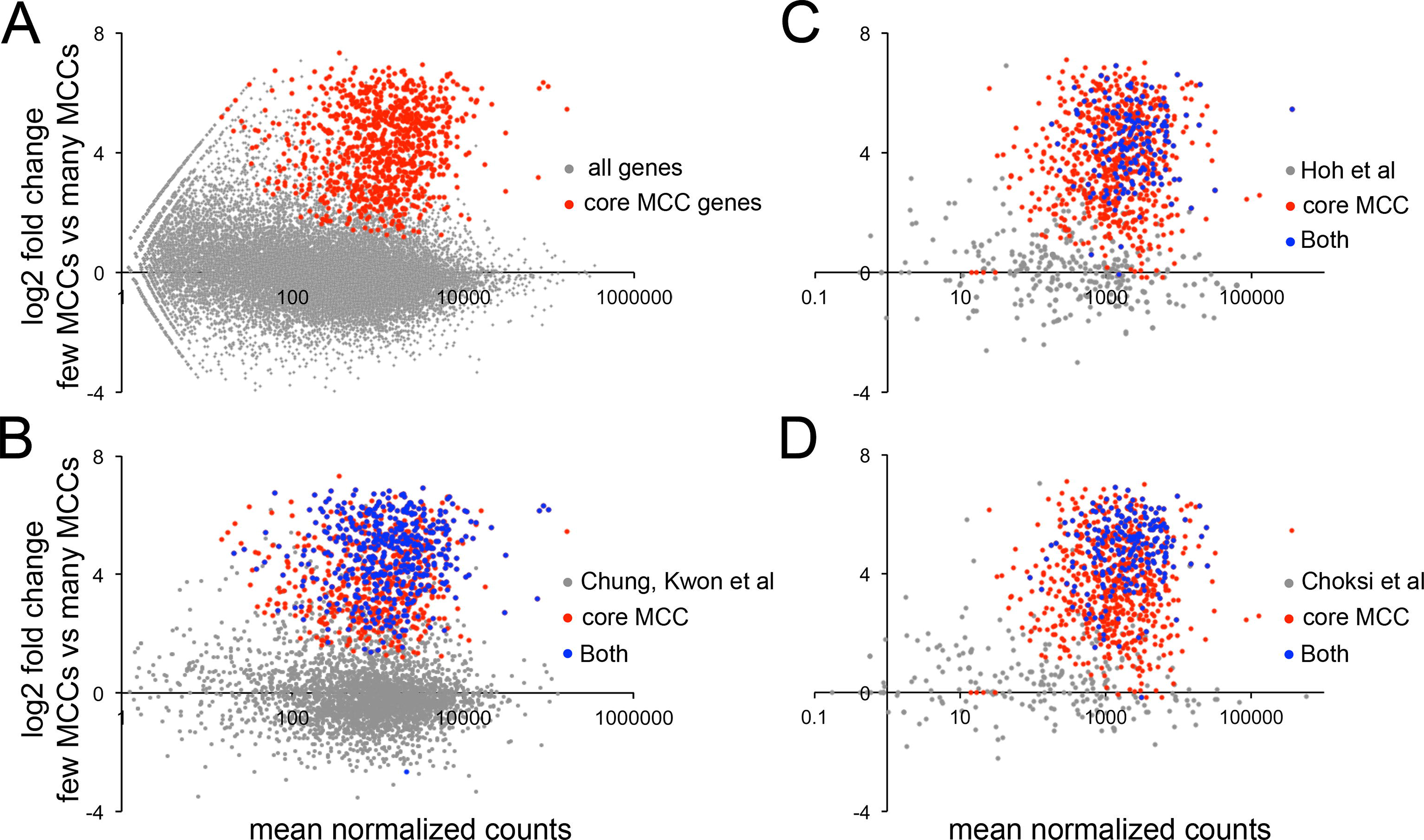
Comparison of motile cilia transcriptomes. **(A)** MA plot (log ratios vs. mean average) of all genes expressed in dissected *X. laevis* ectoderm at the 9 hour timepoint. X axis is normalized counts per gene and the Y axis is log2-fold change of each gene in a comparison between progenitors injected with Notch-icd ("few MCCs") versus progenitors injected with Notch-icd and Multicilin ("many MCCs"). Core MCC genes labeled in red as determined in Fig. 1C, all other *X. laevis* genes labeled in gray. **(B)** Data from (A) but showing only genes from a comparison between in our core MCC list and a multiliciated cell transcriptome obtained by knocking down Rfx2 in *X. laevis* (Chung et al. 2014). Shown are genes only found in our core MCC list (red), genes only found from (Chung et al. 2014), or genes found in both lists. **(C,D)** Many *X. laevis* genes have two paralogs owing to pseudotetraploidy. In order to compare our data to diploid organisms, we collapsed RNAseq counts from paralogs into a single gene. **(C)** Data from (A) but showing only genes in a comparison between our core MCC list and a motile cilia transcriptome obtained by sorting Foxj1+ cells from mouse tracheal epithelial cultures (Hoh et al. 2012). Shown is a similar comparison as (B) and (C). **(D)** Data from (A) but showing only genes from a comparison between our core MCC list and a motile cilia transcriptome obtained by overexpressing Foxj1 in zebrafish embryos (Choksi et al. 2014). Shown are genes only found in our core MCC list (red), genes only found from (Choksi et al. 2014), or genes found in both lists. Please note that as we have collapsed *X. laevis* L and S forms into a single transcript in (C) and (D), the plots will look slightly different.

**Supplemental figure S3:**
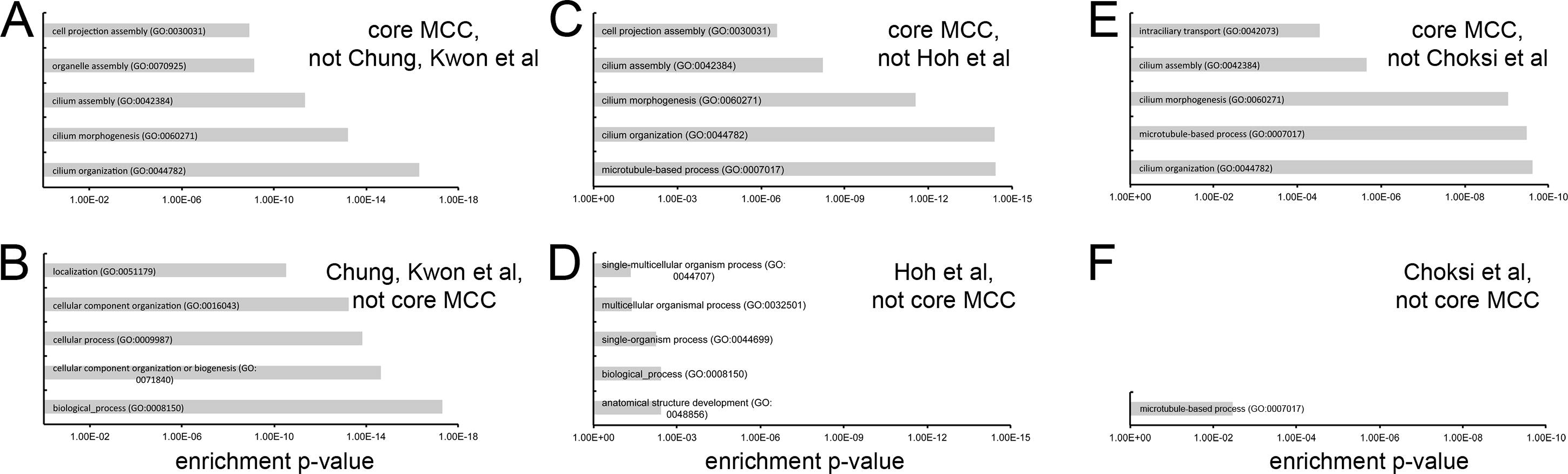
GO term enrichment of core MCC genes versus other lists. **(A,B)** GO term enrichment of genes shown in Supplemental figure S2B that are present on our core MCC list but not regulated by Rfx2 (A), or vice versa (B). **(C,D)** GO term enrichment of genes shown in Supplemental figure S2C that are present on our core MCC list but not expressed in Foxj1+ mouse lung cells (C) or vice versa (D). **(E,F)** GO term enrichment of genes that are shown in Supplemental figure S2D present on our core MCC list but not induced in Zebrafish by Foxj1 overexpression (E) or vice versa (F).

**Supplemental figure S4:**
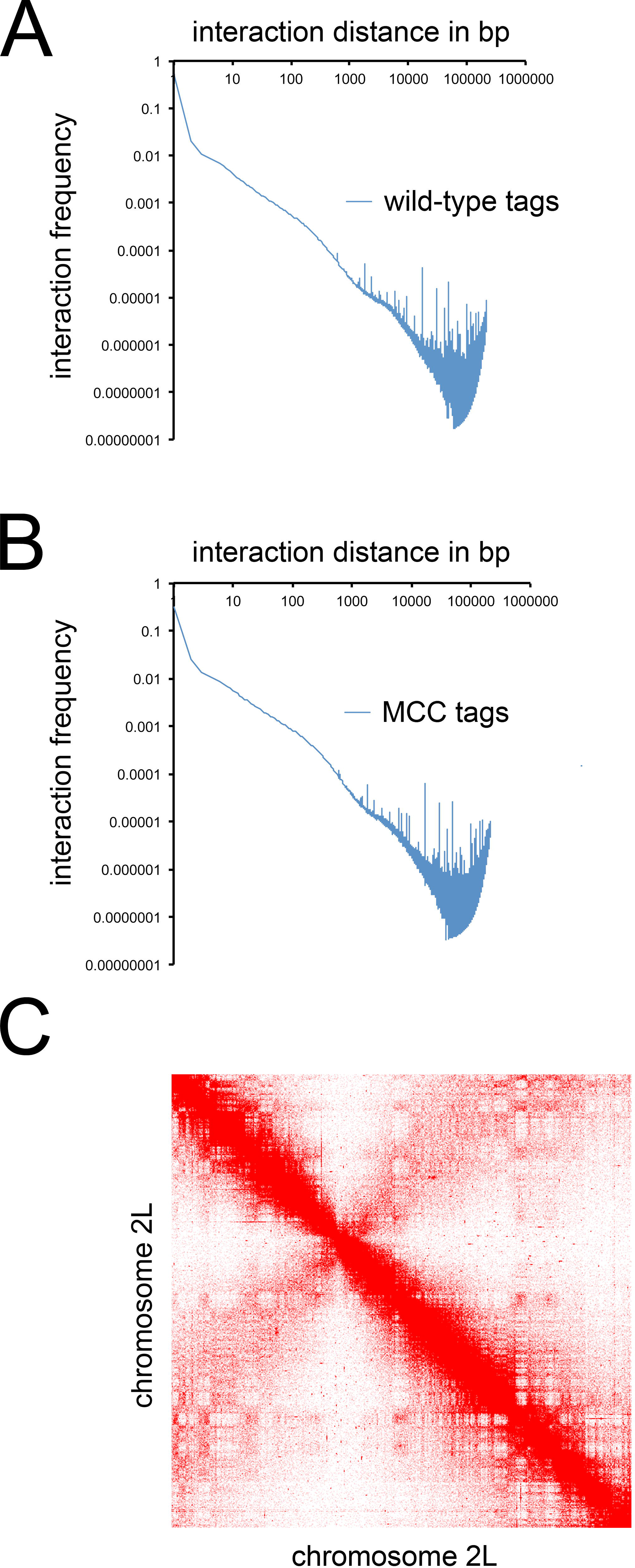
*The 3-dimensional genome structure of* X. laevis *epithelia.* **(A,B)** Tethered conformation capture performed by binding DNA-protein complexes to a fixed substrate followed by digestion by restriction enzymes (in this case, MboI) and proximity ligation. Interaction frequencies are reported to drop as a function of linear genomic distance (Lieberman-Aiden et al. 2009; Lin et al. 2012); the majority of proximity ligation events are the result of immediately neighboring sequence, and not larger 3D structure. Moreover, interchromosomal interactions are thought to be quite rare and strong enrichment for these interactions in the data is suggestive of spurious ligations. We saw high frequencies of local interactions and few interchromosomal interactions in the *X. laevis* genome (A,B). Data are paired-end sequencing tags from a total of 139,003,458 uniquely mapped reads. **(C)** Raw interaction matrix of *X. laevis* chromosome 2L showing contacts across loci.

**Supplemental figure S5:**
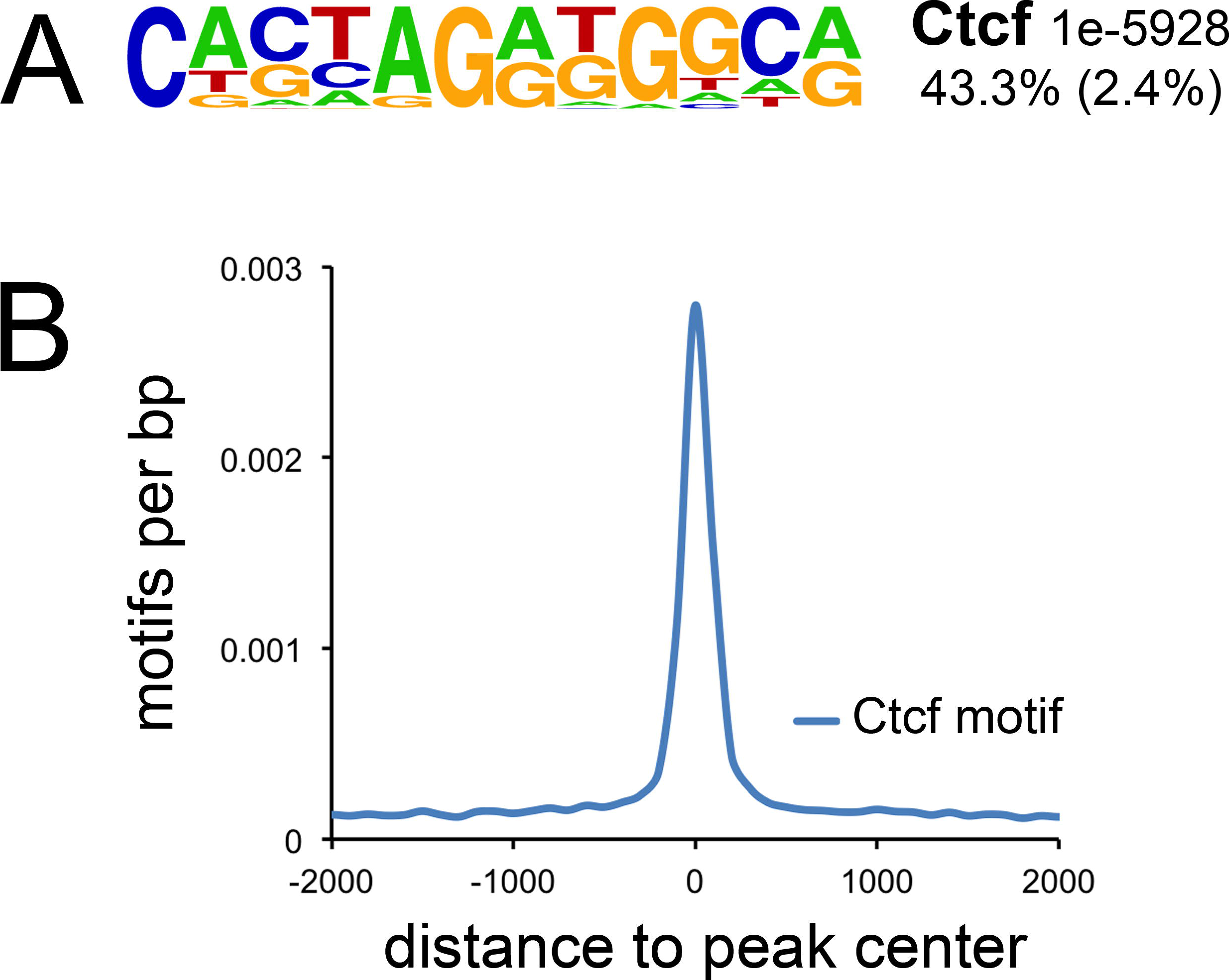
Rad21 ChIPseq confirmation. **(A)** Top *de novo* motif hit from all Rad21 peaks. Top line of label is transcription factor family binding the motif and p-value; second line of label is frequency of motif in peaks versus background frequency (background frequency is in parentheses). **(B)** Frequency and position of top motif in all Rad21 peaks.

**Supplemental figure S6:**
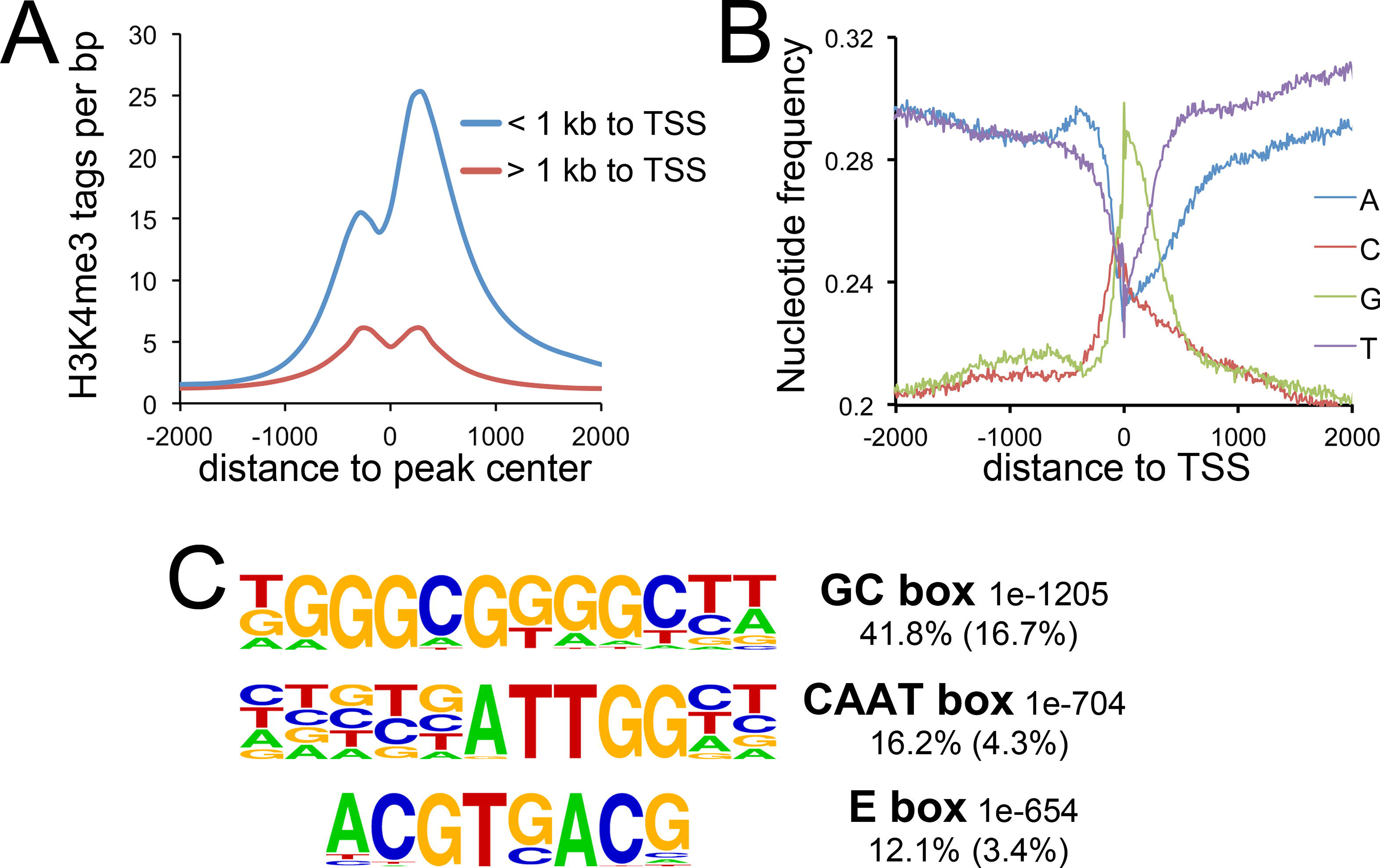
Xenopus laevis *promoter confirmation*. **(A)** Tag counts obtained from H3K4me3 ChIPseq on progenitors overlapping or not overlapping the annotated TSS of *X. laevis* genes (Mayball models, (Chung et al., 2014)). Note dip signifying nucleosome-free region at center of peak. **(B)** Nucleotide frequencies around promoter peak centers. Note GC-rich bias around the TSS (Louie et al., 2011., also see Supplemental figure S10). **(C)** Top *de novo* motifs identified in sequences called as peaks in ChIPseq analysis of all H3K4me3-positive promoters in epithelial progenitors. Top line of label is transcription factor family binding the motif and p-value; second line of label is frequency of motif in peaks versus background frequency (background frequency obtained by searching GC-matched genomic sequence is in parentheses).

**Supplemental figure S7:**
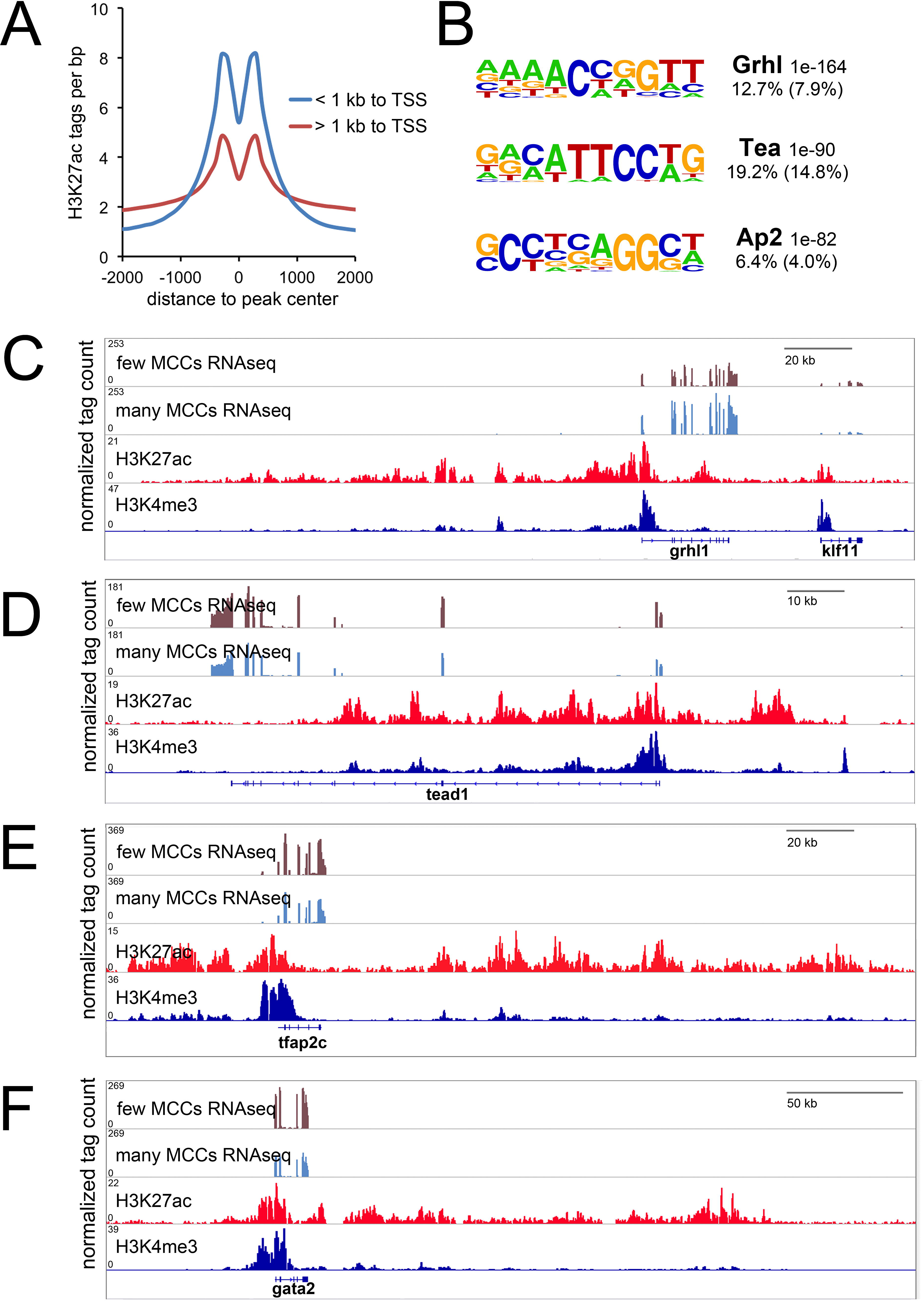
H3K27ac ChIPseq and epithelial superenhancers. **(A)** H3K27ac tag counts around peak centers of confirmed *X*. *laevis* promoters and distal sites obtained by ChIPseq performed on isolated epithelial progenitors. Note nucleosome-free region at dip in center of peak. **(B)** Top *de novo* motif hits from distal H3K27ac peaks. Top line of label is transcription factor family binding the motif and p-value; second line of label is frequency of motif in peaks versus background frequency (background frequency is in parentheses). **(C-F)** Select superenhancers defined by the strategy described in (Whyte et al. 2013). Note superenhancers around key transcription factors **(C-E)** that recognize the most-enriched motifs from all enhancers **(B)**. **(F)** Superenhancer around *gata2*, hinting at an underappreciated role for gata transcription factors in epithelial differentiation.

**Supplemental figure S8:**
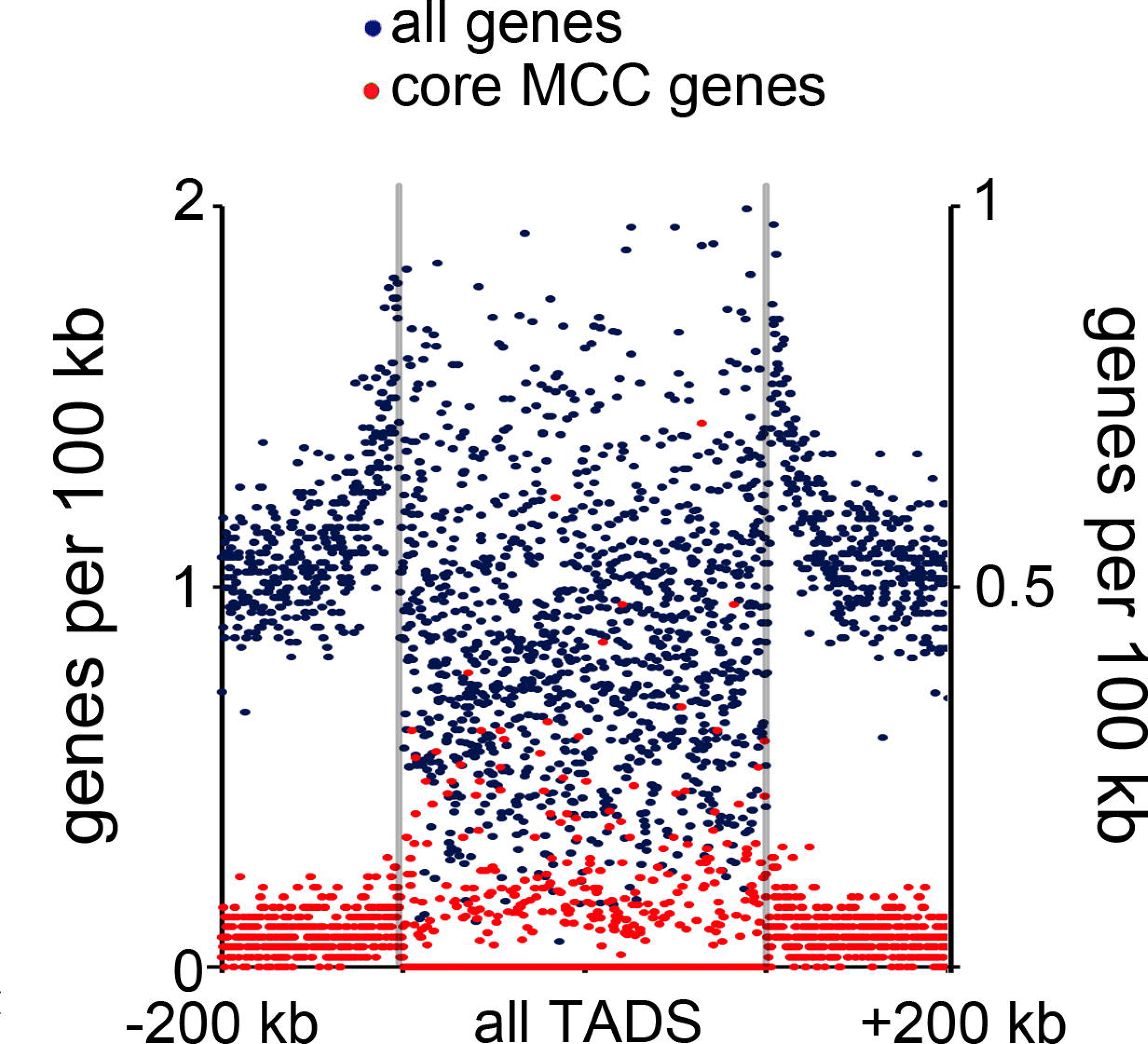
Distribution of core MCC genes in TADs. Metagene plot showing the distribution of core MCC genes relative to all genes in TADs. Note far fewer MCC genes within TADs: bins lacking MCC genes display as dots at bottom of Y axis. Domain region is in the center with boundaries marked, and all domains are normalized to the same size, whereas flanking areas are 200 kb upstream and downstream of those domain boundaries. Each quartile is broken into 175 bins, and each dot denotes one bin.

**Supplemental figure S9:**
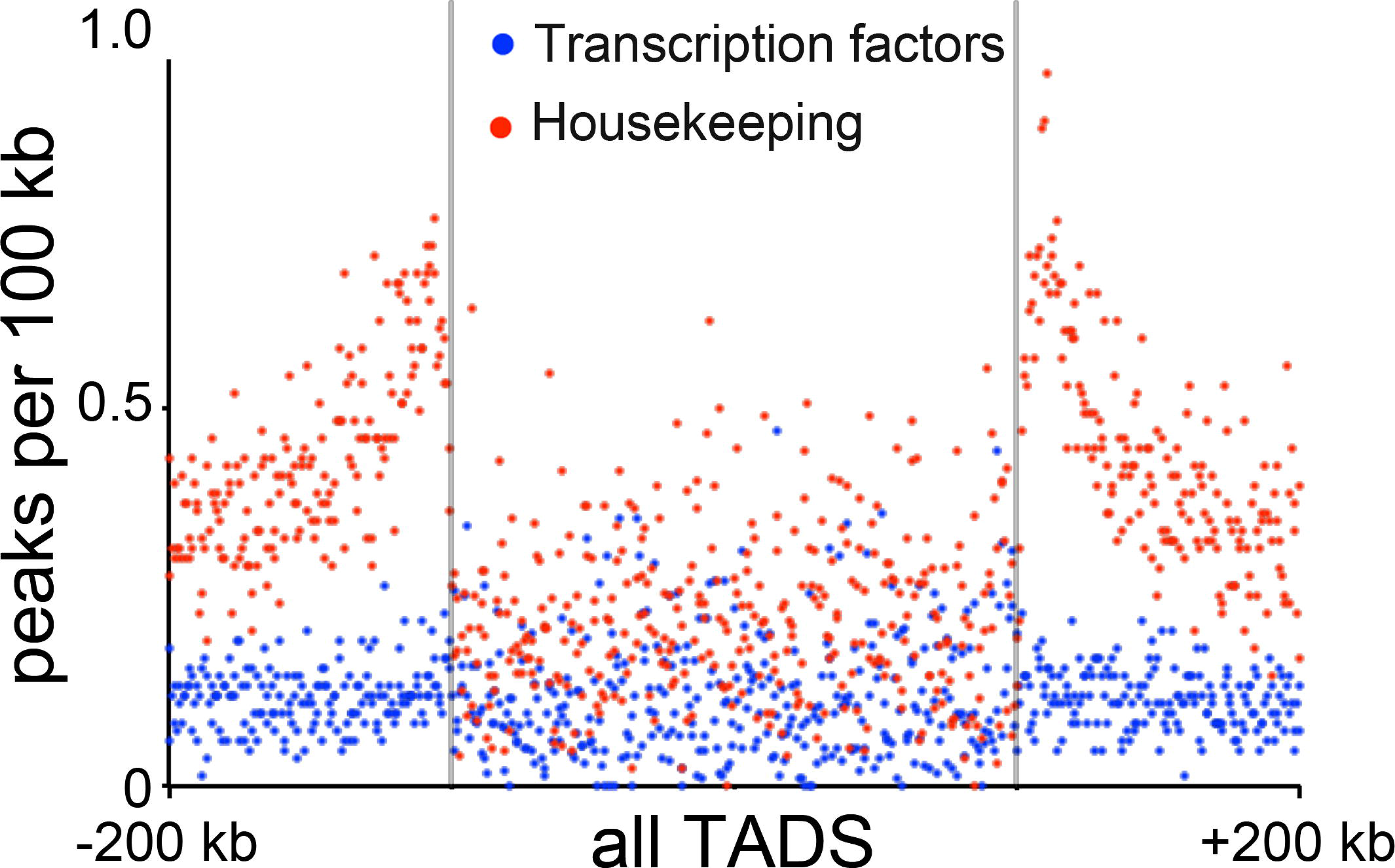
Distribution of transcription factors and housekeeping genes in TADs. Metagene plot showing the distribution of different gene classes relative to all TADs. Domain region is in the center with boundaries marked, and all domains are normalized to the same size, whereas flanking areas are 200 kb upstream and downstream of those domain boundaries. Each quartile is broken into 175 bins, and each dot denotes one bin.

**Supplemental figure S10:**
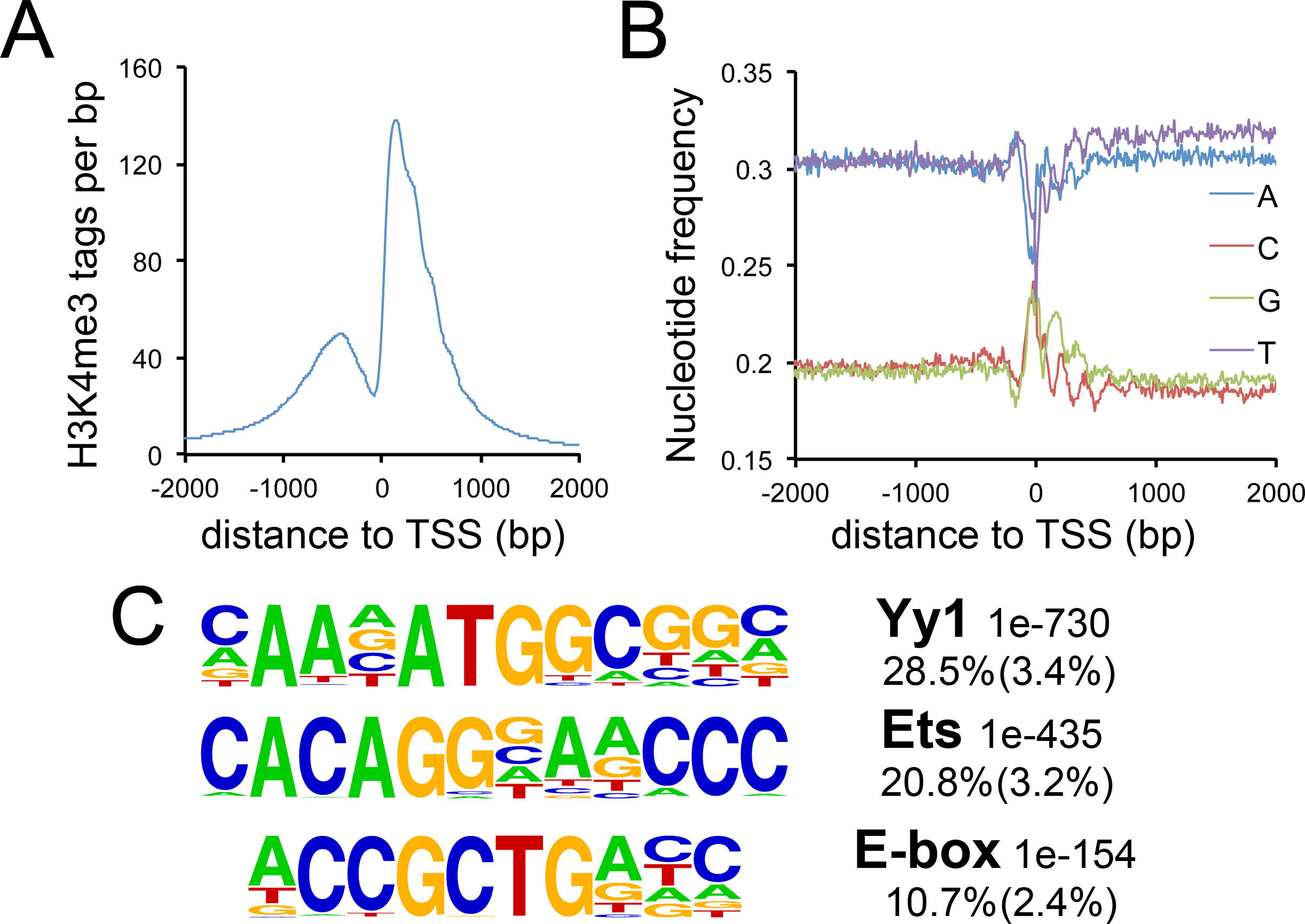
Nematostella vectensis *promoter confirmation*. *N. vectensis* 5’ ends were identified by BLATing (Kent 2002) all 5’ expressed sequence tags to *Nve* genome build v1.0 (Putnam et al. 2007). To confirm these promoters, we then overlapped them with published *N*. *vectensis* H3K4me3 ChIPseq data (Schwaiger et al. 2014). **(A)** Tag counts around peak centers of annotated *Nve* TSS’s. Note nucleosome-free region at center of peak. **(B)** Nucleotide frequencies around promoter peak centers. Note increase in GC-bias around the TSS and nucleosomal periodicity just following. **(C)** Top *de novo* motifs from H3K4me3-positive *N. vectensis* promoters. Note differences in core promoter motif preference relative to *X*. *laevis*, fly, and human (supplemental figure S6, (Louie et al., 2003; Ohler et al, 2002; van Heeringen et al., 2011). Top line of label is transcription factor family binding the motif and p-value; second line of label is frequency of motif in peaks versus background frequency (background frequency is in parentheses).

**Supplemental figure S11:**
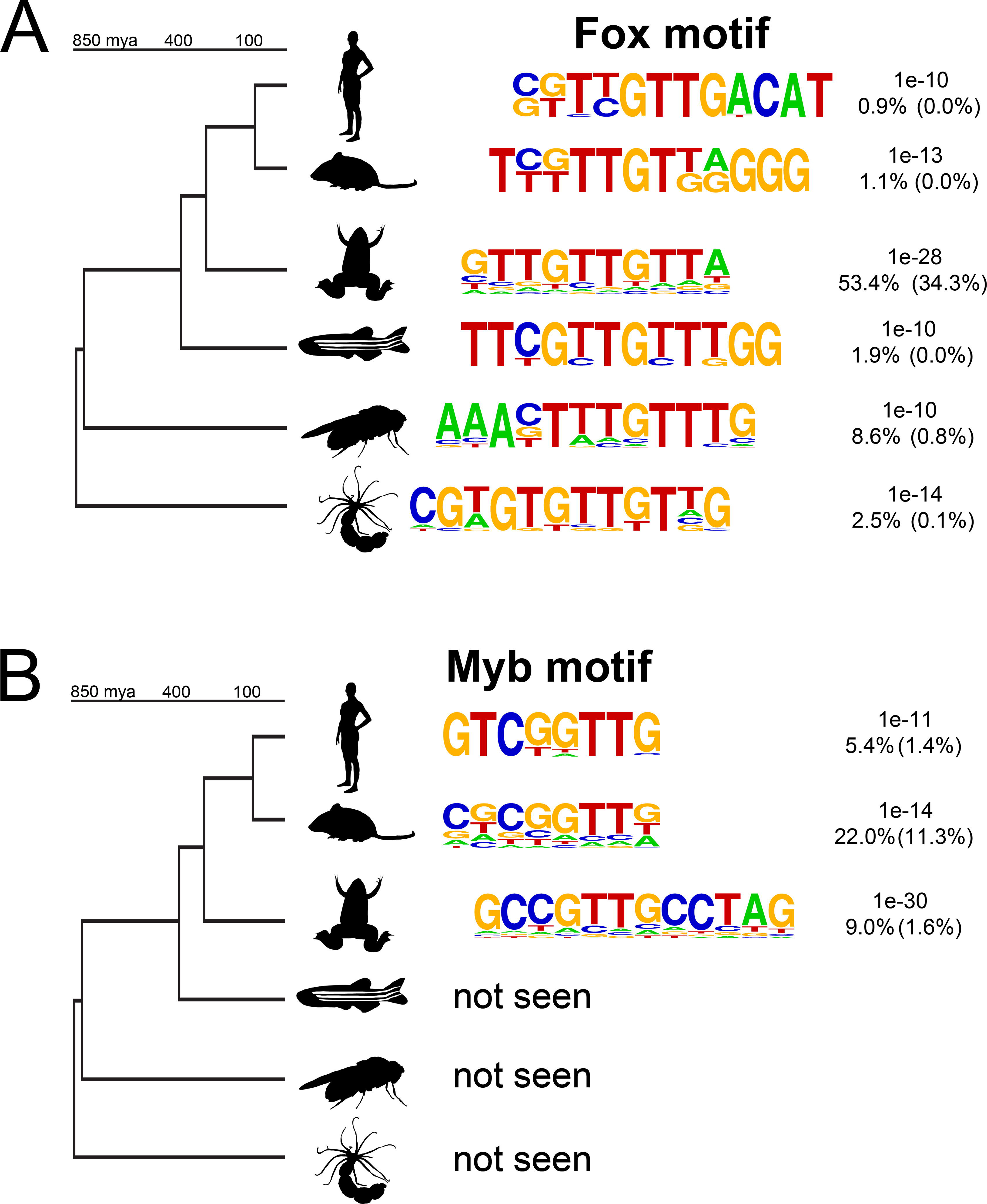
Conservation of MCC promoter architecture. **(A)** Shown are the Forkhead motifs identified *de novo* in the promoters of MCC gene orthologs. Top line of label is p-value; second line of label is frequency of motif in peaks versus background frequency (background frequency is in parentheses). **(B)** Myb motifs in promoters of MCC gene orthologs. Same label convention as in (A).

**Supplemental figure S12:**
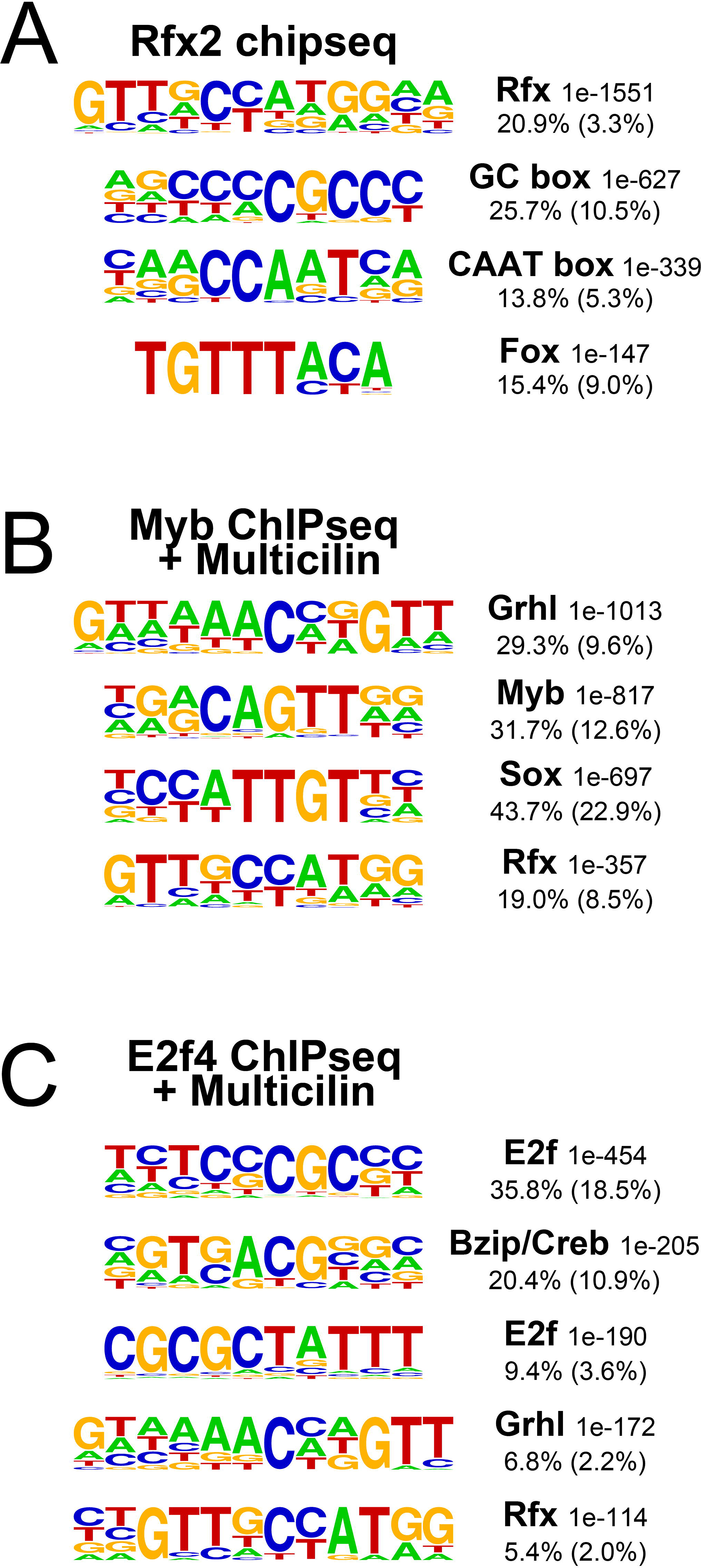
Co-enrichment of motifs in ChIPseq peaks. **(A-C)** Shown are the top *de novo* motifs found enriched in ChIPseq peaks of Rfx2 **(A)**, Myb in the presence of Multicilin **(B)** or E2f4 in the presence of Multicilin **(C)**. Top line of label is transcription factor family binding the motif and p-value; second line of label is frequency of motif in peaks versus background frequency (background frequency is in parentheses). Note overlap of motifs matching factors in these sets (e.g., Rfx motifs in Myb ChIPseq peaks) but reduced representation of these motifs in all promoters (Supplemental figure S6) or all enhancers (Supplemental figure S7).

**Supplemental figure S13:**
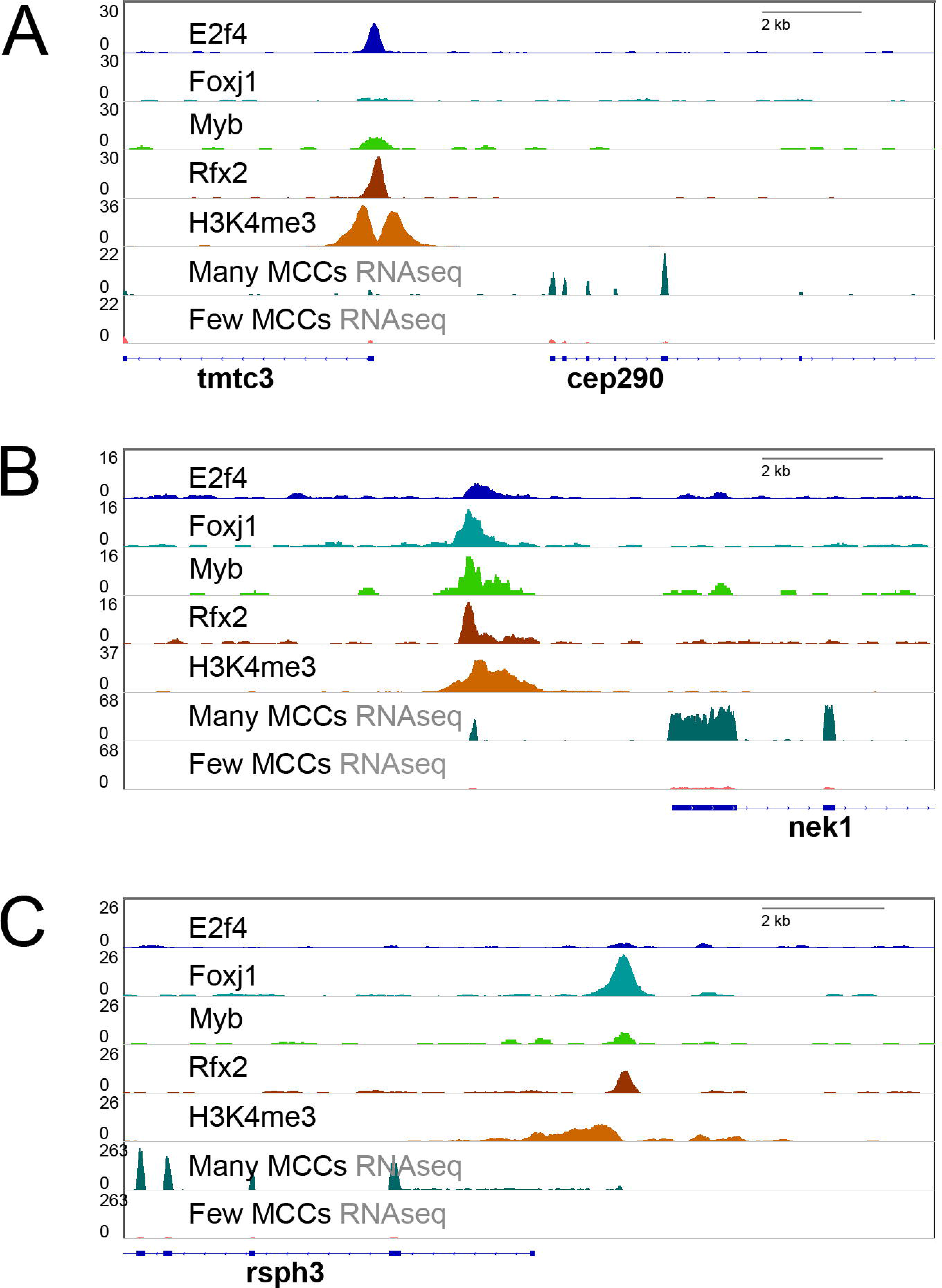
Evidence for misannotated promoters. **(A-C)** Shown are genome browser screenshots of examples of genes with likely promoters outside a 2 kb window centered around transcriptional start sites (TSS’s from the 5’ end of the first exon of the Mayball gene models). Note both enrichment of H3K4me3 and binding of various MCC transcription factors at positions more than 1 kb upstream of described TSS. **(A)** A probable bidirectional promoter; note increased expression of Cep290 in MCCs but no other H3K4me3 peaks nearby. **(B, C)** Note upstream H3K4me3 and transcription factor peaks upstream of 5’-most annotated exons. Also note RNAseq of possible additional upstream exons.

**Supplemental figure S14:**
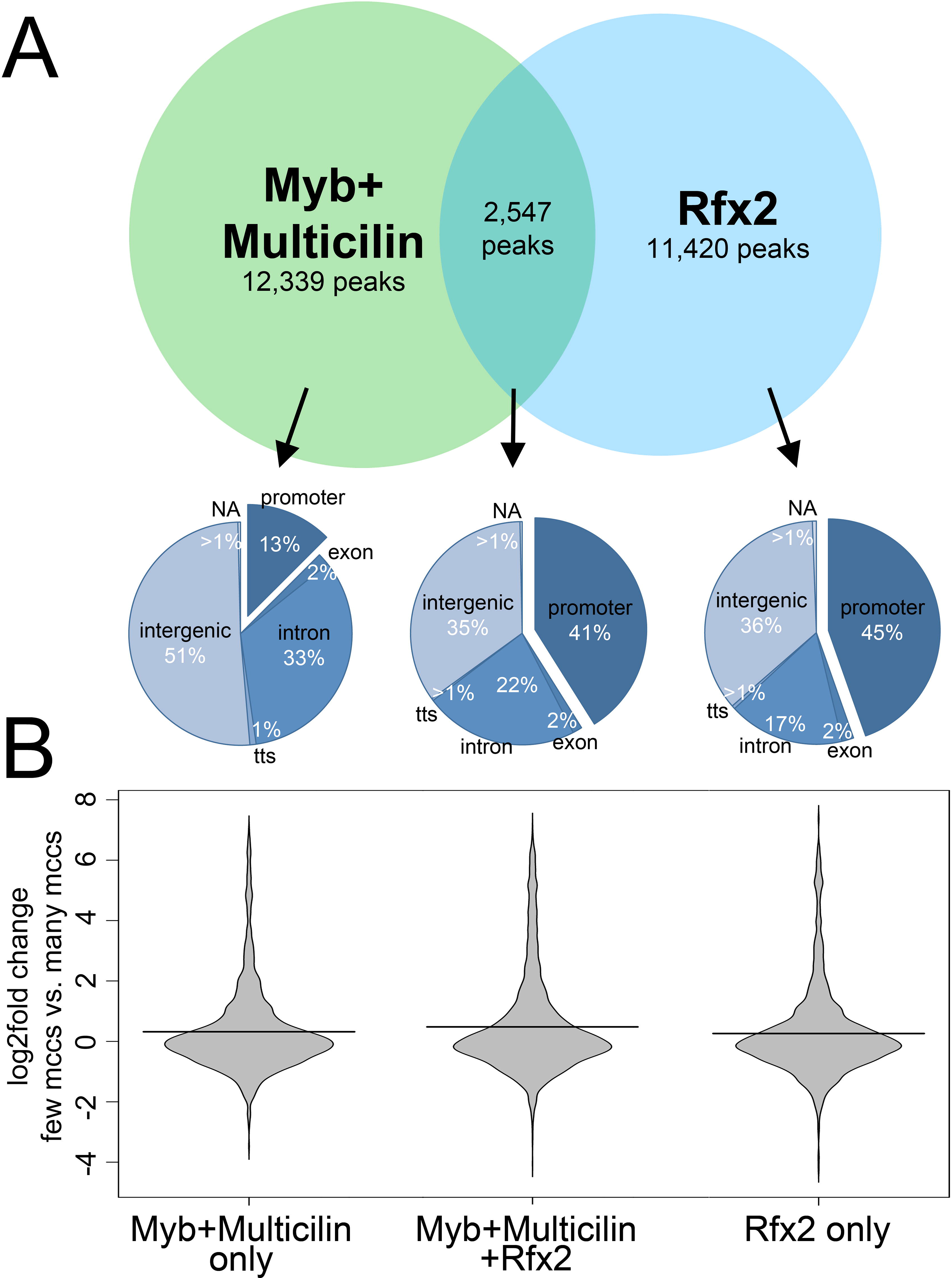
Annotation of Myb ChIPseq peaks in the presence of Multicilin in relation to Rfx2 ChIPseq peaks. **(A)** Sequences were called as peaks using HOMER based on ChIPseq analysis of Rfx2 (Chung et al. 2014) or Myb in the presence of Multicilin. The top Venn diagram represents the overlap of Myb and rRx2 peak sequences, and the piecharts below show the genomic annotations of peaks bound by Myb alone, Rfx2 alone, or both Myb and Rfx2. Promoters are defined as +/- 1kb around the TSS, transcriptional termination sites (TSS) are defined as -100 bp/+1kb around the end of the transcript, and “NA” refers to genomic scaffolds containing no mapped exons. **(B)** Shown is a beanplot summarizing the fold change (log2) in expression at all promoters bound as indicated, based on an RNAseq analysis of progenitor manipulated to repress or promote MCC differentiation (injected with Notch-icd versus Notch-icd and Multicilin).

**Supplemental figure S15:**
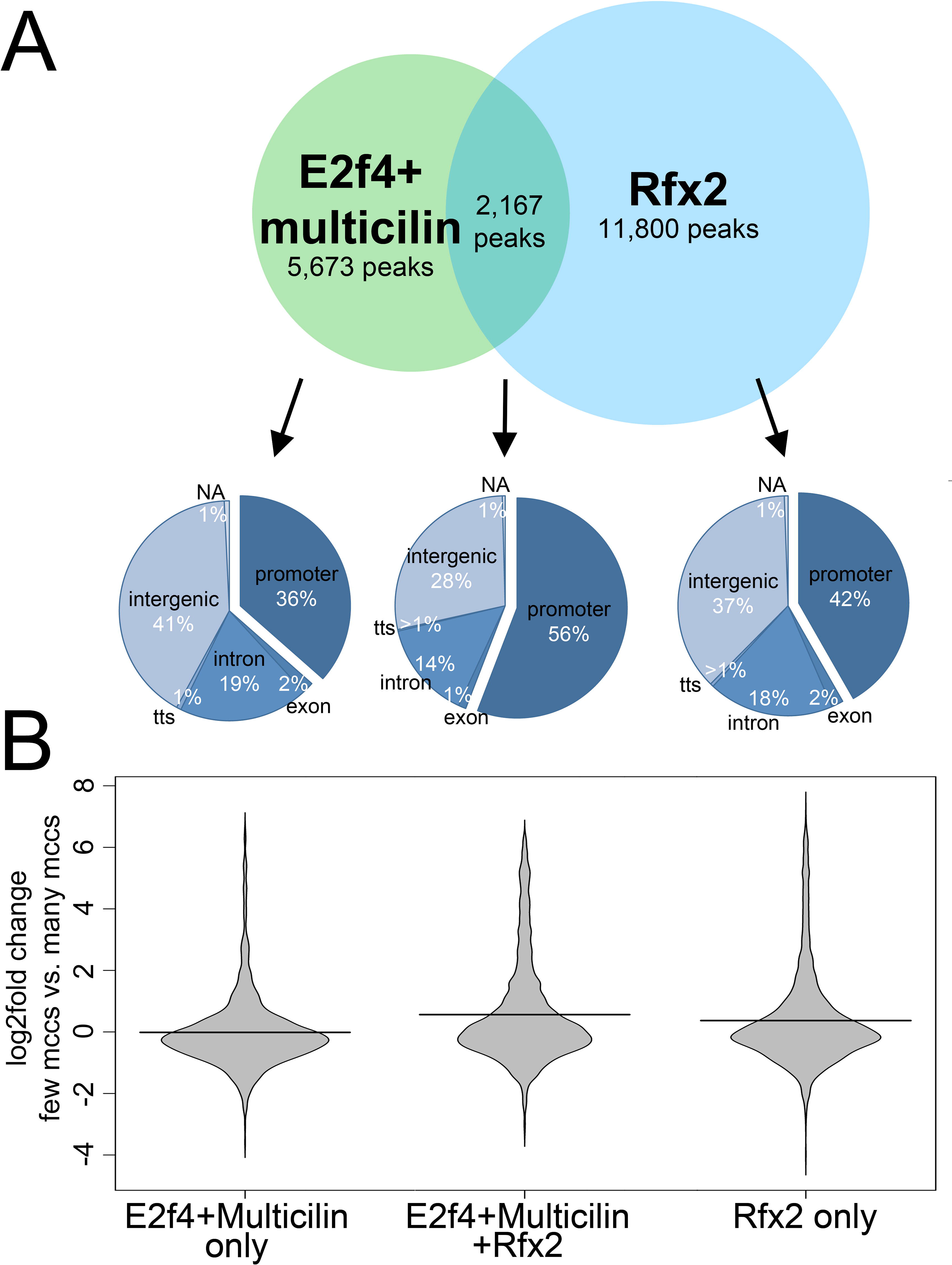
Annotation of E2f4 ChIPseq peaks in the presence of Multicilin in relation to Rfx2 ChIPseq peaks. **(A)** Sequences were called as peaks using HOMER based on ChIPseq analysis of Rfx2 (Chung et al. 2014) or E2f4 in the presence of Multicilin (Ma et al. 2014). The top Venn diagram represents the overlap of E2f4 and Rfx2 peak sequences, and the piecharts below show the genomic annotations of peaks bound by E2f4 alone, Rfx2 alone, or both E2f4 and Rfx2. Promoters are defined as +/- 1kb around the TSS, transcriptional termination sites (TSS) are defined as -100 bp/+1kb around the end of the transcript, and “NA” refers to genomic scaffolds containing no mapped exons. **(B)** Shown is a beanplot summarizing the fold change (log2) in expression at all promoters bound as indicated, based on an RNAseq analysis of progenitors manipulated to repress or promote MCC differentiation (injected with Notch-icd versus Notch-icd and Multicilin).

**Supplemental figure S16:**
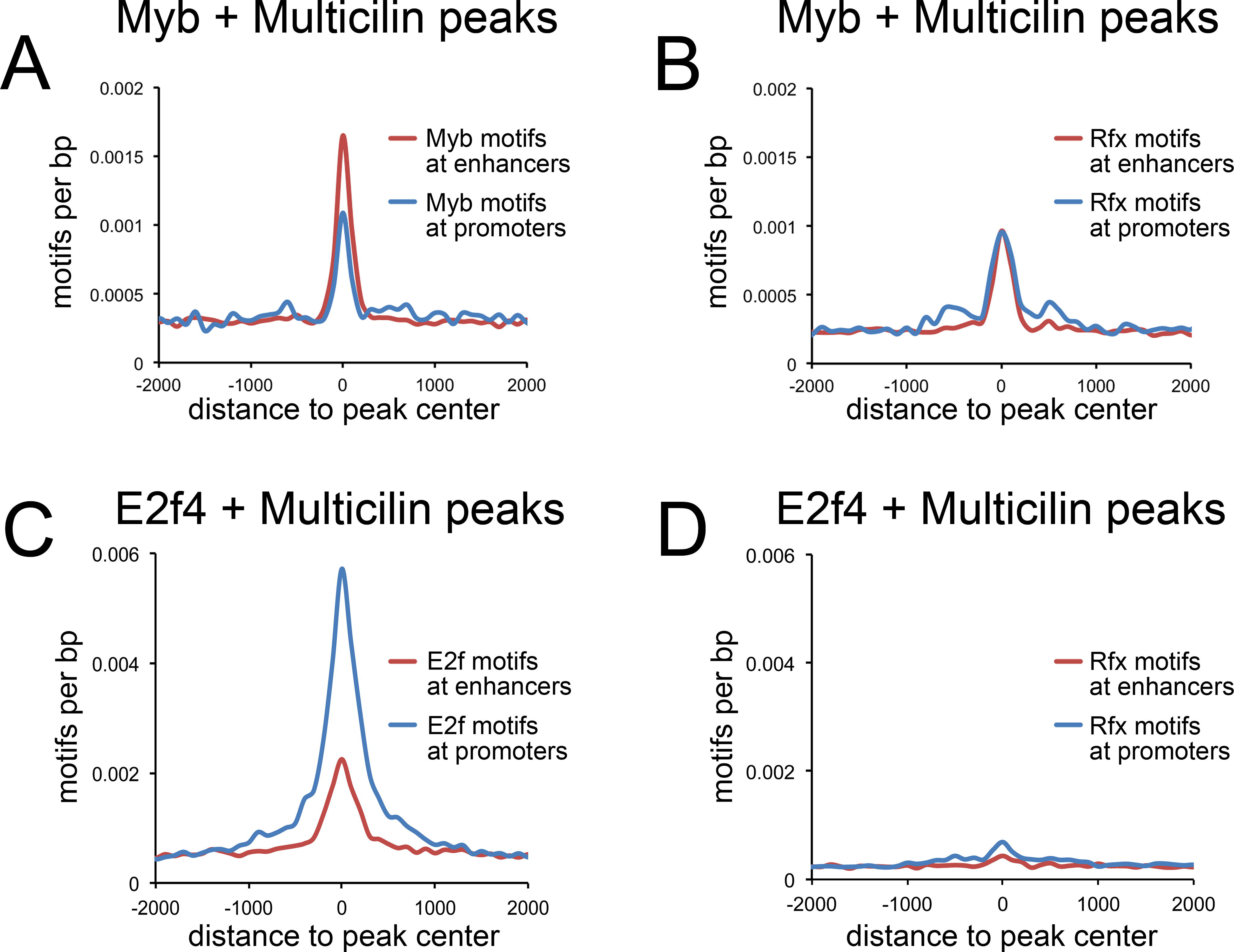
Motif frequency in Myb and E2f4 ChIPseq peaks in the presence of Multicilin. **(A)** Myb motifs in Myb peaks located at promoters and distal sites. **(B)** Rfx motifs in Myb peaks located at promoters and distal sites. **(C)** E2f motifs in E2f4 peaks located at promoters and distal sites. **(D)** Rfx motifs in E2f4 peaks located at promoters and distal sites. Note equal enrichment for Rfx motifs in Myb promoter and distal peaks, similar to Rfx motifs in Foxj1 peaks (Figure 4B).

**Supplemental figure S17:**
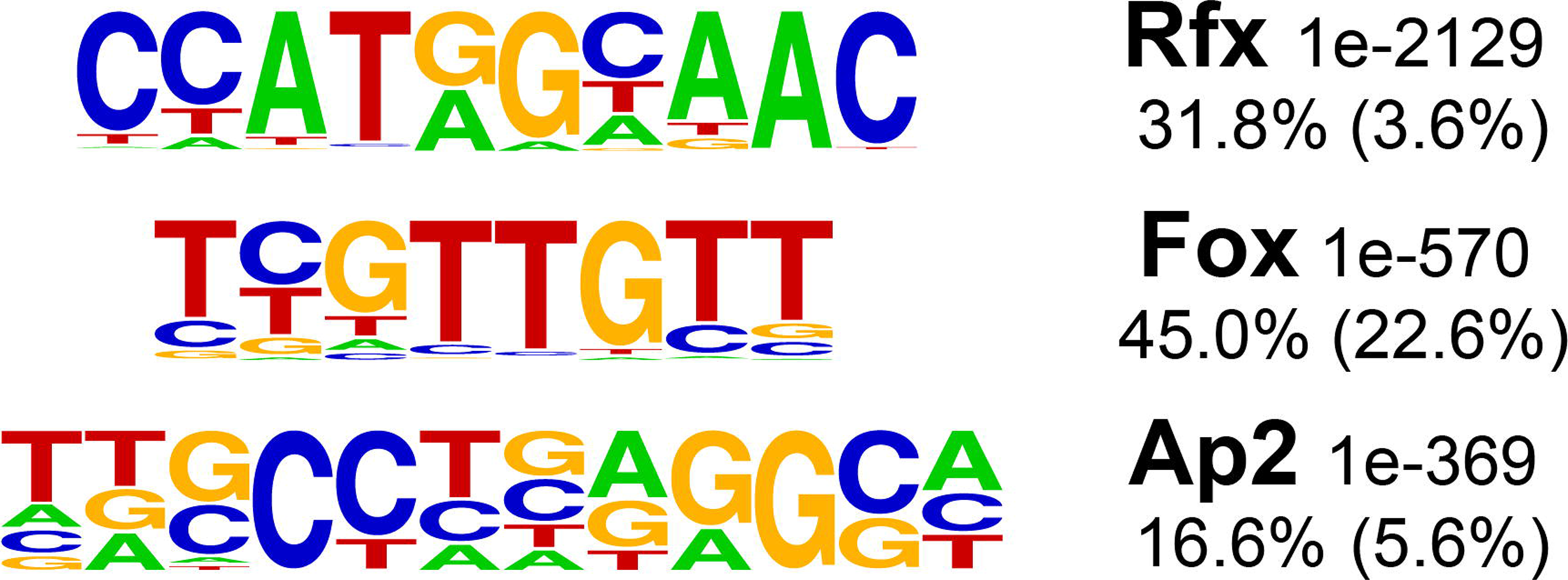
Motif enrichment in Foxj1 ChIPseq peaks not cobound by Rfx2. Shown are the top *de novo* motifs found enriched in ChIPseq peaks of Foxj1 peaks not overlapping with Rfx2 peaks. Top line of label is transcription factor family binding the motif and p-value; second line of label is frequency of motif in peaks versus background frequency (background frequency is in parentheses).

**Supplemental figure S18:**
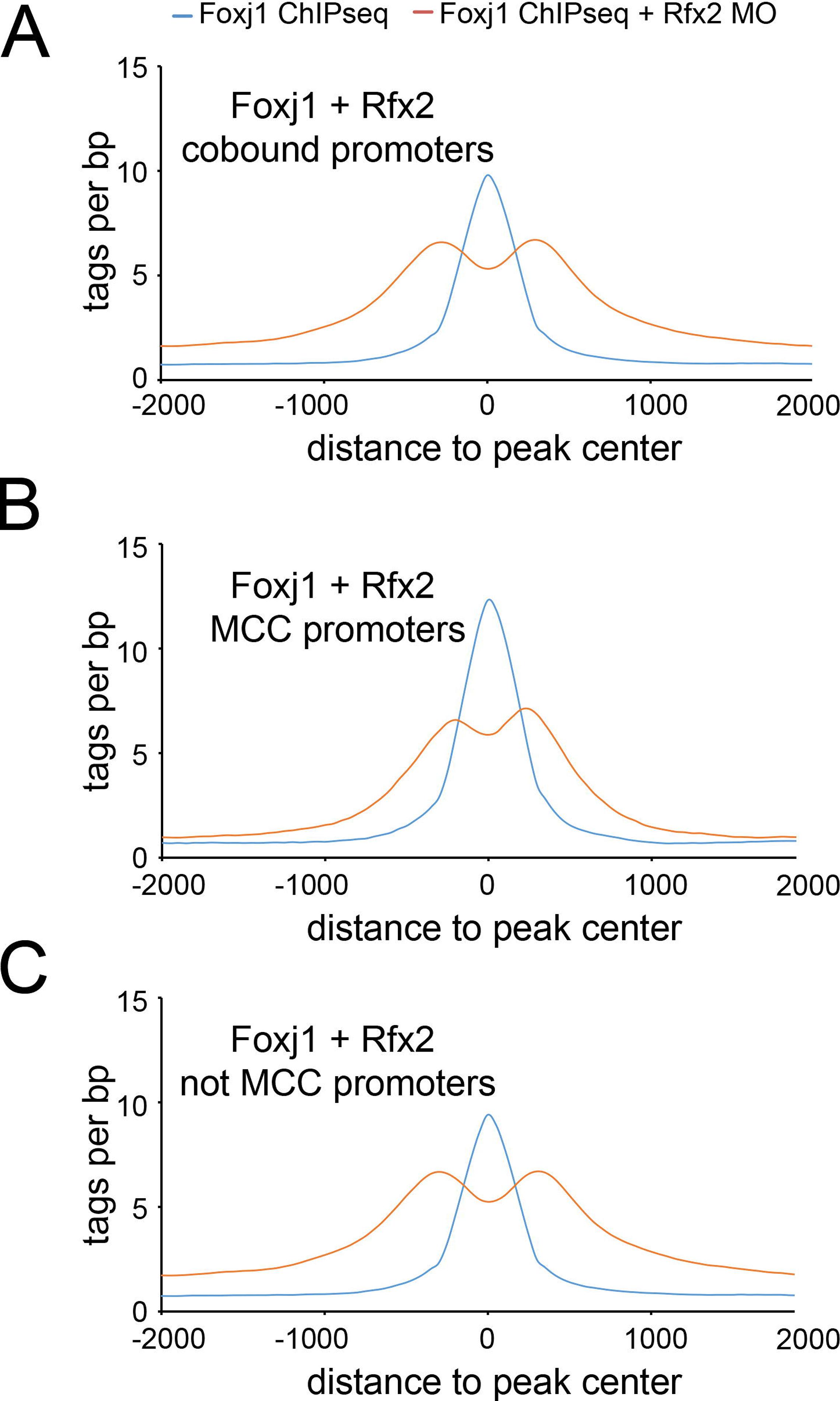
Rfx2 stabilizes Foxj1 more at cobound MCC promoters. Shown are tag counts from Foxj1 ChIPseq and Foxj1 ChIPseq with Rfx2 mopholino at **(A)** all promoters cobound by Foxj1 and Rfx2; **(B)** core MCC promoters cobound by Foxj1 and Rfx2, and **(C)** non-MCC promoters cobound by Foxj1 and Rfx2.

**Supplemental figure S19:**
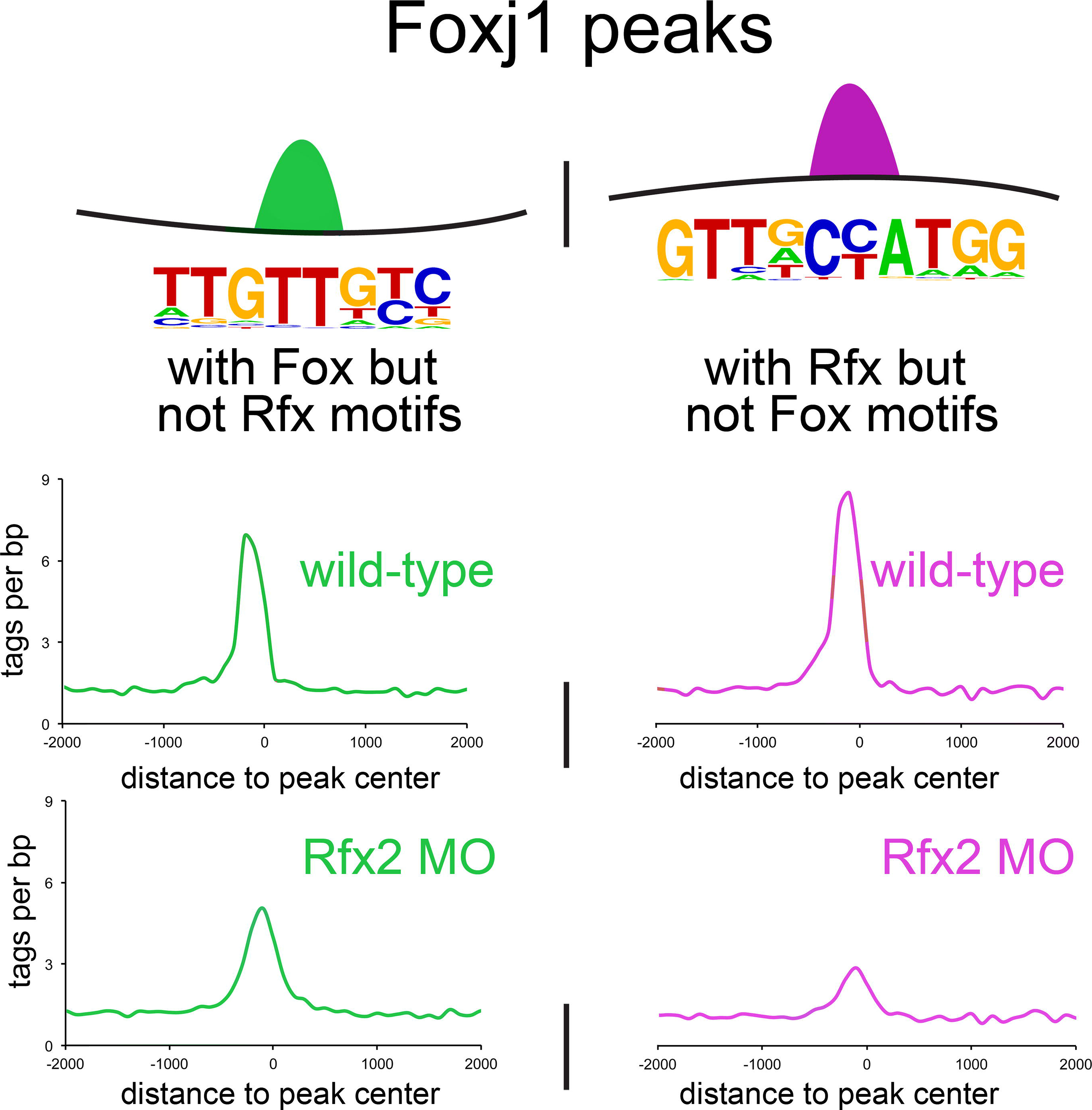
Rfx2 stabilizes Foxj1 more at sites with poor Fox motifs. Shown are tag counts from Foxj1 ChIPseq and Foxj1 ChIPseq with Rfx2 mopholino at genomic positions with a strong Fox mot but no Rfx motif, or a strong Rfx motif and no Fox motif.

## Quigley and Kintner, supplemental tables

Table S1: Manipulations changing cell fates for RNAseq studies

Each manipulation and its effect on MCCs and ionocytes is depicted; cartoons illustrating these manipulations are shown in Figure 1C.

Table S2: *Genes associated with progenitors that become MCCs*

RNAseq analysis was carried out on ectodermal progenitors manipulated by increasing (Notch+) or decreasing (Notch-) signaling, by increasing (Multicilin) or decreasing (dnMulticilin) Multicilin activity, by increasing Foxi1 activity, or by inhibiting E2f4 (dnE2f4) activity. Samples were collected at 3, 6 or 9 hour timepoints, corresponding to the equivalent of stages 13, 16, and 18, respectively, and hierarchical clustering performed on both samples and expression. Listed is a group of genes clustered by similarity of expression as shown in Figure 1F; also see Supplemental Figure S1. A more conservative list, based on the intersection of three comparisons using differential expression and corrected for multiple testing, can be found in Table S7.

Table S3: *Genes associated with progenitors that become ionocytes*

Listed is a group of genes clustered by similarity of expression as determined in Figure S1. A more conservative list, based on differential expression and corrected for multiple testing, can be found in Table S10.

Table S4: *Genes associated with progenitors with increased notch signaling*

Listed is a group of genes clustered by similarity of expression as determined in Figure S2.

Table S5: *Genes associated with progenitors progenitors harvested early*

Listed is a group of genes clustered by similarity of expression as determined in Figure S3.

Table S6: *Genes associated with progenitors harvested late*

Listed is a group of genes clustered by similarity of expression as determined in Figure S4.

Table S7: *Multiciliated cell genes at the 9 hour timepoint*

Top tab of spreadsheet is a group of genes resulting from the intersection of 3 tests for RNAseq differential expression for samples harvested at the 9 hour timepoint (Notch+ vs. Notch, Notch+ vs. Notch+ and Multicilin, and Notch- vs. Notch- and dnMulticilin, see Figure 1E). Additional tabs are results of differential expression testing for all genes across the three comparisons.

Table S8: *Multiciliated cell genes at the 6 hour timepoint*

Top tab of spreadsheet is a group of genes resulting from the intersection of 3 tests for RNAseq differential expression for samples harvested at the 6 hour timepoint (Notch+ vs. Notch, Notch+ vs. Notch+ and Multicilin, and Notch- vs. Notch- and dnMulticilin, see Figure 1E). Additional tabs are results of differential expression testing for all genes across the three comparisons.

Table S9: *Multiciliated cell genes at the 3 hour timepoint*

Top tab of spreadsheet is a group of genes resulting from the intersection of 3 tests for RNAseq differential expression for samples harvested at the 9 hour timepoint (Notch+ vs. Notch, Notch+ vs. Notch+ and Multicilin, and Notch- vs. Notch- and dnMulticilin, see Figure 1E). Additional tabs are results of differential expression testing for all genes across the three comparisons; final tab is intersection between 2 conditions, as intersection of all 3 was sparse.

Table S10: *Ionocyte genes at the 9 hour timepoint*

Listed is a group of genes resulting from differential expression between Notch+ and Notch+ and Foxi1 at the 9 hour timepoint.

Table S11: *Overlap between core MCC genes from this study and those in* Chung, Kwon et al.

Listed, in three tables, are genes found in common between these two datasets and genes unique to either. As both studies derive from *X. laevis* RNAseq, a comparison was made using the Mayball gene model annotations.

Table S12: *Overlap between core MCC genes from this study and those in* Choksi et al.

Listed, in three tables, are genes found in common between these two datasets and genes unique to either. As these data derived from *X. laevis* and *D. rerio,* we converted genes from both studies to human Ensembl annotations to facilitate cross-species comparisons.

Table S13: *Overlap between core MCC genes from this study and those in* Hoh et al.

Listed, in three tables, are genes found in common between these two datasets and genes unique to either. As these data derived from *X. laevis* and *M. musculus,* we converted genes from both studies to human Ensembl annotations to facilitate cross-species comparisons.

Table S14: *Correlation coefficient differences*

Listed is a group of genes found in genomics regins with poor correlation coefficients in 3D conformation data between wild-type and Multicilin-injected epithelial progenitors.

Table S15: *Flanking enhancers per gene*

Listed are all genes and the numbers of active peaks, as determined by H3K4me3 and H3K27ac, that are closer to that gene than any other gene.

Table S16: *Annotated superenhancers*

Listed are the positions and identities of genes nearest to called superenhancers.

Table S17: *List of sequencing experiments*

Listed are details of the various sequencing experiments in this manuscript.

## References

1. Cirillo, L. A. et al. Binding of the winged-helix transcription factor HNF3 to a linker histone site on the nucleosome. EMBO J. 17, 244–54 (1998).

2. Gualdi, R. et al. Hepatic specification of the gut endoderm in vitro: cell signaling and transcriptional control. Genes Dev. 10, 1670–82 (1996).

3. Heinz, S. et al. Simple combinations of lineage-determining transcription factors prime cis-regulatory elements required for macrophage and B cell identities. Mol. Cell 38, 576–89 (2010).

4. Lupien, M. et al. FoxA1 Translates Epigenetic Signatures into Enhancer-Driven Lineage-Specific Transcription. Cell 132, 958–970 (2008).

5. Carter, D., Chakalova, L., Osborne, C. S., Dai, Y. & Fraser, P. Long-range chromatin regulatory interactions in vivo. Nat. Genet. 32, 623–6 (2002).

6. Jin, F. et al. A high-resolution map of the three-dimensional chromatin interactome in human cells. Nature 503, 290–4 (2013).

7. Dixon, J. R. et al. Chromatin architecture reorganization during stem cell differentiation. Nature 518, 331–336 (2015).

8. Lieberman-Aiden, E. et al. Comprehensive mapping of long-range interactions reveals folding principles of the human genome. Science 326, 289–93 (2009).

9. Rao, S. S. P. et al. A 3D Map of the Human Genome at Kilobase Resolution Reveals Principles of Chromatin Looping. Cell 159, 1665–80 (2014).

10. de Laat, W. & Duboule, D. Topology of mammalian developmental enhancers and their regulatory landscapes. Nature 502, 499–506 (2013).

11. Luna-Zurita, L. et al. Complex Interdependence Regulates Heterotypic Transcription Factor Distribution and Coordinates Cardiogenesis. Cell 164, 999–1014 (2016).

12. Choksi, S. P., Lauter, G., Swoboda, P. & Roy, S. Switching on cilia: transcriptional networks regulating ciliogenesis. Development 141, 1427–41 (2014).

13. Brooks, E. R. & Wallingford, J. B. Multiciliated cells. Curr. Biol. 24, R973–82 (2014).

14. Didon, L. et al. RFX3 modulation of FOXJ1 regulation of cilia genes in the human airway epithelium. Respir. Res. 14, 70 (2013).

15. Newton, F. G. et al. Forkhead transcription factor Fd3F cooperates with Rfx to regulate a gene expression program for mechanosensory cilia specialization. Dev. Cell 22, 1221–33 (2012).

16. Lister, R. et al. Highly integrated single-base resolution maps of the epigenome in Arabidopsis. Cell 133, 523–36 (2008).

17. Wettstein, D. A., Turner, D. L. & Kintner, C. The Xenopus homolog of Drosophila Suppressor of Hairless mediates Notch signaling during primary neurogenesis. Development 124, 693–702 (1997).

18. Deblandre, G. A., Wettstein, D. A., Koyano-Nakagawa, N. & Kintner, C. A two-step mechanism generates the spacing pattern of the ciliated cells in the skin of Xenopus embryos. Development 126, 4715–28 (1999).

19. Stubbs, J. L., Vladar, E. K., Axelrod, J. D. & Kintner, C. Multicilin promotes centriole assembly and ciliogenesis during multiciliate cell differentiation. Nat. Cell Biol. 14, 140–7 (2012).

20. Choksi, S. P., Babu, D., Lau, D., Yu, X. & Roy, S. Systematic discovery of novel ciliary genes through functional genomics in the zebrafish. Development 141, 3410–9 (2014).

21. Hoh, R. A., Stowe, T. R., Turk, E. & Stearns, T. Transcriptional program of ciliated epithelial cells reveals new cilium and centrosome components and links to human disease. PLoS One 7, e52166 (2012).

22. Chung, M.-I. et al. Coordinated genomic control of ciliogenesis and cell movement by RFX2. Elife 3, e01439 (2014).

23. Kalhor, R., Tjong, H., Jayathilaka, N., Alber, F. & Chen, L. Genome architectures revealed by tethered chromosome conformation capture and population-based modeling. Nat. Biotechnol. 30, 90–8 (2012).

24. Nagano, T. et al. Comparison of Hi-C results using in-solution versus in-nucleus ligation. Genome Biol. 16, 175 (2015).

25. Lin, Y. C. et al. Global changes in the nuclear positioning of genes and intra- and interdomain genomic interactions that orchestrate B cell fate. Nat. Immunol. 13, 1196–204 (2012).

26. Dixon, J. R. et al. Topological domains in mammalian genomes identified by analysis of chromatin interactions. Nature 485, 376–80 (2012).

27. Andrey, G. et al. A switch between topological domains underlies HoxD genes collinearity in mouse limbs. Science 340, 1234167 (2013).

28. Zuin, J. et al. Cohesin and CTCF differentially affect chromatin architecture and gene expression in human cells. Proc. Natl. Acad. Sci. 111, 996–1001 (2013).

29. Sofueva, S. et al. Cohesin-mediated interactions organize chromosomal domain architecture. EMBO J. 32, 3119–29 (2013).

30. Heinz, S., Romanoski, C. E., Benner, C. & Glass, C. K. The selection and function of cell type-specific enhancers. Nat. Rev. Mol. Cell Biol. 16, 144–54 (2015).

31. van Heeringen, S. J. et al. Nucleotide composition-linked divergence of vertebrate core promoter architecture. Genome Res. 21, 410–21 (2011).

32. Zhang, H.-M. et al. AnimalTFDB: a comprehensive animal transcription factor database. Nucleic Acids Res. 40, D144–9 (2012).

33. Eisenberg, E. & Levanon, E. Y. Human housekeeping genes, revisited. Trends Genet. 29, 569–74 (2013).

34. Elkon, R. et al. RFX transcription factors are essential for hearing in mice. Nat. Commun. 6, 8549 (2015).

35. Piasecki, B. P., Burghoorn, J. & Swoboda, P. Regulatory Factor X (RFX)-mediated transcriptional rewiring of ciliary genes in animals. Proc. Natl. Acad. Sci. U. S. A. 107, 12969–74 (2010).

36. Lenhard, B., Sandelin, A. & Carninci, P. Metazoan promoters: emerging characteristics and insights into transcriptional regulation. Nat. Rev. Genet. 13, 233–45 (2012).

37. Louie, E., Ott, J. & Majewski, J. Nucleotide frequency variation across human genes. Genome Res. 13, 2594–601 (2003).

38. Ohler, U., Liao, G., Niemann, H. & Rubin, G. M. Computational analysis of core promoters in the Drosophila genome. Genome Biol. 3, RESEARCH0087 (2002).

39. Stubbs, J. L., Oishi, I., Izpisúa Belmonte, J. C. & Kintner, C. The forkhead protein Foxj1 specifies node-like cilia in Xenopus and zebrafish embryos. Nat. Genet. 40, 1454–60 (2008).

40. Ma, L., Quigley, I., Omran, H. & Kintner, C. Multicilin drives centriole biogenesis via E2f proteins. Genes Dev. 28, 1461–1471 (2014).

41. Tan, F. E. et al. Myb promotes centriole amplification and later steps of the multiciliogenesis program. Development 140, 4277–86 (2013).

42. Session, A. M. et al. Genome evolution in the allotetraploid frog Xenopus laevis. Nature 538, 336–343 (2016).

43. Heintzman, N. D. et al. Histone modifications at human enhancers reflect global cell-type-specific gene expression. Nature 459, 108–12 (2009).

44. Yao, L. et al. Inferring regulatory element landscapes and transcription factor networks from cancer methylomes. Genome Biol. 16, 105 (2015).

45. Nemajerova, A. et al. TAp73 is a central transcriptional regulator of airway multiciliogenesis. Genes Dev. (2016). doi: 10.1101/gad.279836.116

46. Efimenko, E. et al. Analysis of xbx genes in C. elegans. Development 132, 1923–34 (2005).

47. Vij, S. et al. Evolutionarily ancient association of the FoxJ1 transcription factor with the motile ciliogenic program. PLoS Genet. 8, e1003019 (2012).

48. Chung, M.-I. et al. RFX2 is broadly required for ciliogenesis during vertebrate development. Dev. Biol. 363, 155–65 (2012).

49. Ernst, J. & Kellis, M. Interplay between chromatin state, regulator binding, and regulatory motifs in six human cell types. Genome Res. 23, 1142–54 (2013).

50. John, S. et al. Chromatin accessibility pre-determines glucocorticoid receptor binding patterns. Nat. Genet. 43, 264–8 (2011).

51. Katan-Khaykovich, Y. & Shaul, Y. RFX1, a Single DNA-binding Protein with a Split Dimerization Domain, Generates Alternative Complexes. J. Biol. Chem. 273, 24504–24512 (1998).

52. Reith, W. et al. MHC class II regulatory factor RFX has a novel DNA-binding domain and a functionally independent dimerization domain. Genes Dev. 4, 1528–40 (1990).

53. Ji, X. et al. 3D Chromosome Regulatory Landscape of Human Pluripotent Cells. Cell Stem Cell 18, 262–275 (2015).

54. Zhu, L. J. et al. ChIPpeakAnno: a Bioconductor package to annotate ChIP-seq and ChIP-chip data. BMC Bioinformatics 11, 237 (2010).

55. Geremek, M. et al. Gene expression studies in cells from primary ciliary dyskinesia patients identify 208 potential ciliary genes. Hum. Genet. 129, 283–93 (2011).

56. Shawlot, W., Vazquez-Chantada, M., Wallingford, J. B. & Finnell, R. H. Rfx2 is required for spermatogenesis in the mouse. Genesis (2015). doi: 10.1002/dvg.22880

57. Aftab, S., Semenec, L., Chu, J. S.-C. & Chen, N. Identification and characterization of novel human tissue-specific RFX transcription factors. BMC Evol. Biol. 8, 226 (2008).

58. Xie, X. et al. Systematic discovery of regulatory motifs in conserved regions of the human genome, including thousands of CTCF insulator sites. Proc. Natl. Acad. Sci. U. S. A. 104, 7145–50 (2007).

59. Zhang, M., Bolfing, M. F., Knowles, H. J., Karnes, H. & Hackett, B. P. Foxj1 regulates asymmetric gene expression during left-right axis patterning in mice. Biochem. Biophys. Res. Commun. 324, 1413–20 (2004).

60. Garg, A., Futcher, B. & Leatherwood, J. A new transcription factor for mitosis: in Schizosaccharomyces pombe, the RFX transcription factor Sak1 works with forkhead factors to regulate mitotic expression. Nucleic Acids Res. 43, 6874–88 (2015).

61. Banerji, J., Olson, L. & Schaffner, W. A lymphocyte-specific cellular enhancer is located downstream of the joining region in immunoglobulin heavy chain genes. Cell 33, 729–40 (1983).

62. Dostie, J. et al. Chromosome Conformation Capture Carbon Copy (5C): a massively parallel solution for mapping interactions between genomic elements. Genome Res. 16, 1299–309 (2006).

63. Dowen, J. M. et al. Control of Cell Identity Genes Occurs in Insulated Neighborhoods in Mammalian Chromosomes. Cell 159, 374–387 (2014).

64. Tang, Z. et al. CTCF-Mediated Human 3D Genome Architecture Reveals Chromatin Topology for Transcription. Cell 163, 1611–1627 (2015).

65. Duttke, S. H. C. et al. Human promoters are intrinsically directional. Mol. Cell 57, 674–84 (2015).

66. Sive, H. L., Grainger, R. M. & Harland, R. M. Microinjection of Xenopus oocytes. Cold Spring Harb. Protoc. 2010, pdb.prot5536 (2010).

67. Quigley, I. K., Stubbs, J. L. & Kintner, C. Specification of ion transport cells in the Xenopus larval skin. Development 138, 705–14 (2011).

68. Dobin, A. et al. STAR: ultrafast universal RNA-seq aligner. Bioinformatics 29, 15–21 (2013).

69. Roberts, A. & Pachter, L. Streaming fragment assignment for real-time analysis of sequencing experiments. Nat. Methods 10, 71–3 (2013).

70. de Hoon, M. J. L., Imoto, S., Nolan, J. & Miyano, S. Open source clustering software. Bioinformatics 20, 1453–4 (2004).

71. Saldanha, A. J. Java Treeview--extensible visualization of microarray data. Bioinformatics 20, 3246–8 (2004).

72. Anders, S. et al. Count-based differential expression analysis of RNA sequencing data using R and Bioconductor. Nat. Protoc. 8, 1765–86 (2013).

73. Li, H. & Durbin, R. Fast and accurate short read alignment with Burrows-Wheeler transform. Bioinformatics 25, 1754–60 (2009).

74. Thorvaldsdóttir, H., Robinson, J. T. & Mesirov, J. P. Integrative Genomics Viewer (IGV): high-performance genomics data visualization and exploration. Brief. Bioinform. 14, 178–92 (2013).

75. Putnam, N. H. et al. Sea anemone genome reveals ancestral eumetazoan gene repertoire and genomic organization. Science 317, 86–94 (2007).

76. Kent, W. J. BLAT--the BLAST-like alignment tool. Genome Res. 12, 656–64 (2002).

77. Schwaiger, M. et al. Evolutionary conservation of the eumetazoan gene regulatory landscape. Genome Res. 24, 639–50 (2014).

78. Hedges, S. B., Marin, J., Suleski, M., Paymer, M. & Kumar, S. Tree of life reveals clock-like speciation and diversification. Mol. Biol. Evol. 32, 835–45 (2015).

79. Ramsay, R. G. & Gonda, T. J. MYB function in normal and cancer cells. Nat. Rev. Cancer 8, 523–34 (2008).

80. Chai, X., Nagarajan, S., Kim, K., Lee, K. & Choi, J. K. Regulation of the boundaries of accessible chromatin. PLoS Genet. 9, e1003778 (2013).

81. Kalhor, R., Tjong, H., Jayathilaka, N., Alber, F. & Chen, L. Genome architectures revealed by tethered chromosome conformation capture and population-based modeling. Nat. Biotechnol. 30, 90–8 (2012).

82. Shannon, P. et al. Cytoscape: a software environment for integrated models of biomolecular interaction networks. Genome Res. 13, 2498–504 (2003).

